# Circulating chromatin reveals the effects of disease-associated variants on gene regulation

**DOI:** 10.1101/2025.11.12.688010

**Authors:** Ziwei Zhang, Surya B. Chhetri, Karl Semaan, Ze Zhang, Zhenjie Jin, Shahabeddin Sotudian, Liming Liang, Alexander Gusev, Sylvan Baca

## Abstract

A fundamental challenge in human genetics is determining how variation in regulatory DNA shapes complex traits and disease risk. Chromatin quantitative trait loci (cQTLs) can address this challenge by revealing the effects of disease-linked genetic variants on regulatory element activity. Discovering cQTLs in disease-relevant tissues at scale remains challenging, however. To address this limitation, we leveraged advances in epigenomic liquid biopsy. We profiled histone modifications in circulating chromatin from patients with cancer to identify cell-free chromatin QTLs (cfcQTLs). By sampling cancer-derived chromatin in plasma, we captured cfcQTLs affecting regulatory elements from diverse non-hematologic tissues, as well as developmentally restricted elements that are reactivated in cancer (enriched 16-fold). Applying a cistrome-wide association study (CWAS), we linked 4,891 cfcQTLs to 1,011 traits and diseases. Developmentally restricted cfcQTLs that were not found in white blood cells were associated with 22.7 traits per 100 QTLs, compared to 0.58 for WBC-restricted cQTLs, underscoring the power of cfcQTLs for capturing genetic variation that shapes phenotypes. We extended our approach beyond germline variants to non-coding somatic mutations in cancer by measuring the activating effects of *TERT* promoter mutations from circulating chromatin. This work provides a path to scalable discovery of cQTLs across tissues, diseases, and populations to dissect the genetics of common diseases, enabled by the ease of sampling blood.

## Introduction

Many common genetic variants contribute to human traits and disease risk through effects on non-coding regulatory DNA. Noncoding variants account for a substantial proportion of phenotype heritability, often exceeding the contribution from protein-coding variants to the genetic architecture of complex traits and diseases^1–3^. Genome-wide association studies (GWAS) have identified thousands of disease risk loci in regulatory DNA, where genetic variants likely affect gene expression by altering the activity of regulatory elements. Yet, it remains a fundamental challenge to determine how non-coding genetic variants at these loci confer disease risk^4–6^. Unlike protein-coding variants, no “code” exists to predict the molecular consequences of regulatory variants on chromatin. A further challenge is that genetic effects on chromatin or gene regulation that may be context-dependent, manifesting only in certain cell types or states^7–10^, making them difficult to measure in differentiated, bulk tissues at steady-state.

Expression quantitative trait loci (eQTLs) studies have helped decode the regulatory effects of genetic variants by correlating genotypes with gene expression levels^11,12^. However, eQTLs identified in bulk tissues or easily accessible cell types (mainly peripheral blood cells) often fail to capture disease-relevant genetic effects, as many risk variants operate in specific cellular contexts that may be rare, transient, or difficult to sample^13–16^. Recent estimates suggest that known cis-eQTLs mediate only ∼11% of trait heritability on average, leaving a substantial proportion of disease-associated genetic variation unexplained^17,18^.

Chromatin QTLs (cQTLs) are emerging as a powerful tool for characterizing the functional impacts of genetic variation in non-coding DNA. cQTLs are genetic variants whose genotype correlates with quantitative measurements of a nearby regulatory element, such as chromatin accessibility, transcription factor binding, or histone modifications^8,19,20^. Measuring genetic effects on chromatin state through cQTL profiling can provide insights that extend beyond those offered by eQTLs^21^. cQTLs explain a higher proportion of heritability for complex traits compared to eQTLs, suggesting they may capture a broader spectrum of functional effects that are missed by studying gene expression alone^8,10,21–23^. Analysis of cQTLs can aid in identifying disease-causing variants and understanding how they function. For example, cQTLs can reveal risk variants that disrupt nucleotide motifs bound by specific transcription factors controlling the expression of nearby genes^8,24–27^. Coupled with techniques for linking regulatory elements to the genes they control^8,28–30^, cQTL analysis can identify causal steps in the pathway from a genetic variant to cancer development – an ultimate goal of genetics.

A major bottleneck in cQTL discovery is the limited scalability of current approaches. Identifying cQTLs requires epigenomic profiling across large cohorts. For this reason, the vast majority of cQTL discovery has been performed only in peripheral blood cells, because they are easily accessible compared to other tissues^31,32^. However, due to the context-dependent nature of gene regulation, peripheral blood cQTLs only capture a fraction of gene regulatory elements and fail to capture many tissue-specific regulatory elements.

Cell-free DNA (cfDNA) circulating in blood offers a promising approach for addressing these scalability challenges. Because cfDNA is derived from diverse tissues throughout the body, it provides a window into regulatory landscapes of non-hematologic tissues that may otherwise be inaccessible^33,34^. In a wide variety of human conditions – pregnancy^35^, cancer^36,37^, autoimmune disease^38–40^, ischemia^41,42^, neurodegenerative disease^43,44^, and sepsis^45^ – affected tissues shed DNA into peripheral blood. The first liquid biopsy applications focused on detecting genetic alterations in cancer and fetal aneuploidy in pregnancy and are now in widespread clinical use^46^. Recently, technological advances have enabled measurements of epigenomic profiles from cfDNA^36^. As a result, a broad range of features that correlate with regulatory element activity, such as DNA methylation, histone modifications, or nucleosome positioning, can be measured from blood plasma^47,48^. This provides an opportunity to study the effect of gene regulation on chromatin in disease-relevant tissues, at scale. In this study, we identified cQTLs from circulating chromatin (cfcQTLs) for two histone modifications, H3K4me3 and H3K27ac, demonstrating that cfcQTLs efficiently capture a broad range of trait-linked cQTLs that are not detectable in peripheral blood cells.

## Results

### Circulating cancer chromatin reveals regulatory element activity across diverse tissues

We leveraged a large atlas of circulating cell-free chromatin profiles (N=751) from 366 individuals, comprising 328 cancer patients representing 14 distinct cancer types and 38 healthy controls^48^. This resource includes 303 H3K27ac profiles (marking active enhancers and promoters) and 448 H3K4me3 profiles (predominantly marking active promoters) assessed using cell-free chromatin immunoprecipitation sequencing (cfChIP-seq; **Fig. 1a**). Across patients with cancer, the median estimated circulating tumor DNA (ctDNA) content^37^ was 0.051 (range: 0-0.969; **Fig. 1b**). The specificity of cfChIP signal for capturing regulatory elements was high: 99.0% of H3K27ac cfPeaks (cell-free chromatin peaks, consensus genomic regions with enriched signal across samples) and 98.4% of H3K4me3 cfPeaks directly overlapped ChIP-seq peaks observed in Roadmap^49^ (**Extended Fig. 1a**).

**Fig. 1:**
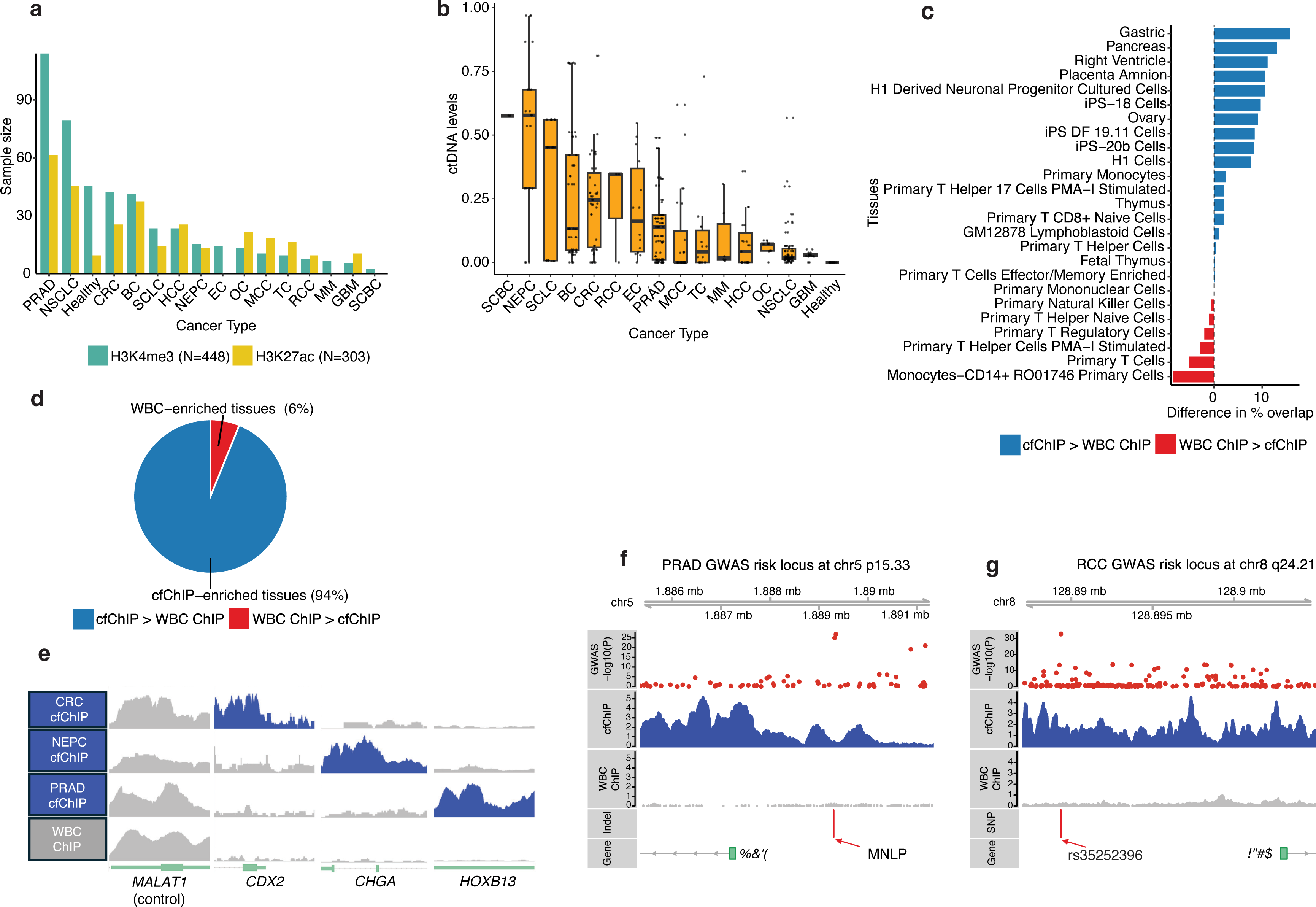
Circulating chromatin reveals regulatory element activity in non-hematologic tissues. **(a)** cfChIP-seq profiles for two histone modifications across 15 cancer types and healthy individuals. Cancer types shown (left to right): PRAD, prostate adenocarcinoma; NSCLC, non-small cell lung cancer; Melanoma; CRC, colorectal cancer; SCLC, small cell lung cancer; LCCC, large cell lung carcinoma; HCC, hepatocellular carcinoma; HETCC, hepatocellular carcinoma; EC, esophageal cancer; GC, gastric cancer; MCC, Merkel cell carcinoma; TC, thymic carcinoma; RCC, renal cell carcinoma; MM, multiple myeloma; GBM, glioblastoma; SCBC, small cell bladder cancer. **(b)** cfDNA tumor fraction across cancer types. **(c)** Differences in regulatory element overlap between cfChIP-seq peaks and WBC ChIP-seq peaks across 98 EpiMap reference epigenomes. Positive values (blue) indicate greater cfChIP overlap; negative values (red) indicate greater WBC ChIP overlap. **(d)** Proportion of reference epigenomes with more overlapping cfChIP peaks (blue, 94%) versus more overlapping WBC ChIP-seq peaks (red, 6%). **(e)** Normalized H3K27ac cfChIP-seq and WBC ChIP-seq signal at selected genes with lineage-restricted expression. Tracks show representative samples from colorectal cancer (CRC), neuroendocrine prostate cancer (NEPC), prostate adenocarcinoma (PRAD), and white blood cells (WBC). **(f)** Normalized H3K27ac signals and GWAS association p-values at a prostate cancer risk locus 5p15.33. A validated multiple-nucleotide-length polymorphism (MNLP) at this locus is highlighted. **(g)** Normalized H3K27ac cfChIP-seq signals at a renal cancer risk locus 8q24.21. A causal variant (rs35252396) is highlighted that affects HIF binding and expression of MYC and PVT1.

We hypothesized that circulating chromatin from patients with cancer captures a broad array of tissue-specific regulatory elements that are not observable in chromatin from peripheral blood cells. We compared cfPeaks and peaks derived from H3K27ac ChIP-seq of peripheral blood cell populations, including CD14+ monocytes (n=162), CD16+ neutrophils (n=174), and naive CD4+ T cells (n=142) from the BLUEPRINT Project^32,50^. We observed substantial enrichment of cfPeaks for overlap with non-hematologic tissues compared to WBC and depletion of overlap with leukocyte epigenomes (**Fig. 1c**). Overall, cfPeaks exhibited higher percentages of overlap than WBC peaks in 92 of 98 reference epigenomes^49^ (**Fig. 1d, Supplementary Table 1**), with greater capture of tissue-restricted regulatory elements **(Supplemental Note)**. Consistent with this broader tissue representation, cfChIP-seq signals were enriched at lineage-restricted genes such as *CDX2*^51–53^ in colorectal cancer, *CHGA*^54^ in neuroendocrine prostate cancer, and *HOXB13*^55–57^ in prostate adenocarcinoma, while these signals were nearly absent in WBC ChIP-seq (**Fig. 1e**). These results demonstrate that cfChIP-seq captures regulatory elements from non-hematologic tissues that extend beyond the repertoire accessible from peripheral blood cells.

Intriguingly, cfChIP-seq identified cell-type-specific regulatory element signals at loci implicated in disease risk by GWAS. For instance, we observed a high H3K27ac signal in circulating chromatin from prostate cancer patients at the p15.33 prostate cancer risk locus^58–63^, where a multiple nucleotide length polymorphism (MNLP) increases androgen receptor (AR) binding ^26^, boosting oncogenic transcription of *IRX4* (**Fig. 1f**). Another example is the renal cell carcinoma (RCC) GWAS risk locus at *8q24.21*, where SNP rs35252396 increases binding of hypoxia-inducible transcription factors (HIFs), which is necessary for *MYC* and *PVT1* expression in cells with renal tubular origin^64^ (**Fig. 1g**). Importantly, H3K27ac signals at *p15.33* and *8q24.21* were strong in cfChIP-seq data but essentially undetectable with WBC ChIP-seq. These data support the possibility of measuring tissue-specific regulatory element activity at disease risk loci using cfChIP-seq, which we turned to next.

### Circulating chromatin reveals genetic effects on regulatory element activity

Building on our observation that cfChIP-seq captures regulatory element signals from diverse tissues, we asked whether genetic determinants of enhancer and promoter activity could be identified from the cfChIP-seq data. We imputed SNP genotypes directly from cfChIP-seq reads^8^ and analyzed *cis*-genetic effects on chromatin modifications using two complementary approaches: assessing allelic imbalance (AI) at heterozygous SNPs^24,65^ and identifying chromatin quantitative trait loci (cQTL; **Fig. 2a**)^8,66^. We implemented a combined test for identifying SNPs with either AI or cQTLs, which we denote as cfcQTL SNPs. Using this approach, we identified 11,834 H3K27ac and 18,160 H3K4me3 cfcQTLs, corresponding to 4,790 H3K27ac peaks (9.28% of total) and 7,361 H3K4me3 peaks (14.26% of total) with significant cfcQTLs, which we term cfcPeaks (**Fig. 2b**). Among the identified cfcQTLs, 9 H3K4me3 and 12 H3K27ac cfcQTL variants were recently reported to affect gene expression across multiple cell types^67^ and associate with cancer risk^68–73^ (**Supplementary Table 3**). For example, rs60143196, located within an H3K4me3 peak on chr3, exhibited significant allelic imbalance in our cfChIP-seq dataset (AF = 0.35, P = 9.54 × 10⁻¹⁶) with nearly 2-fold higher read counts for the reference allele compared to the alternative allele (**Fig. 2c**). Similarly, rs7729529 at chr5:43,006,900-43,009,350 showed significant allelic imbalance favoring the alternative allele (AF = 0.55, P = 2.15 × 10⁻⁴), with alternative allele read counts exceeding reference allele counts (**Fig. 2d**). Both SNPs have been linked to lung adenocarcinoma susceptibility^72^ affecting regulatory element activity in primary bronchial and tracheal epithelial cell lines. These results demonstrate that cfcQTLs can identify functionally relevant genetic variants in disease-relevant tissues.

**Fig. 2:**
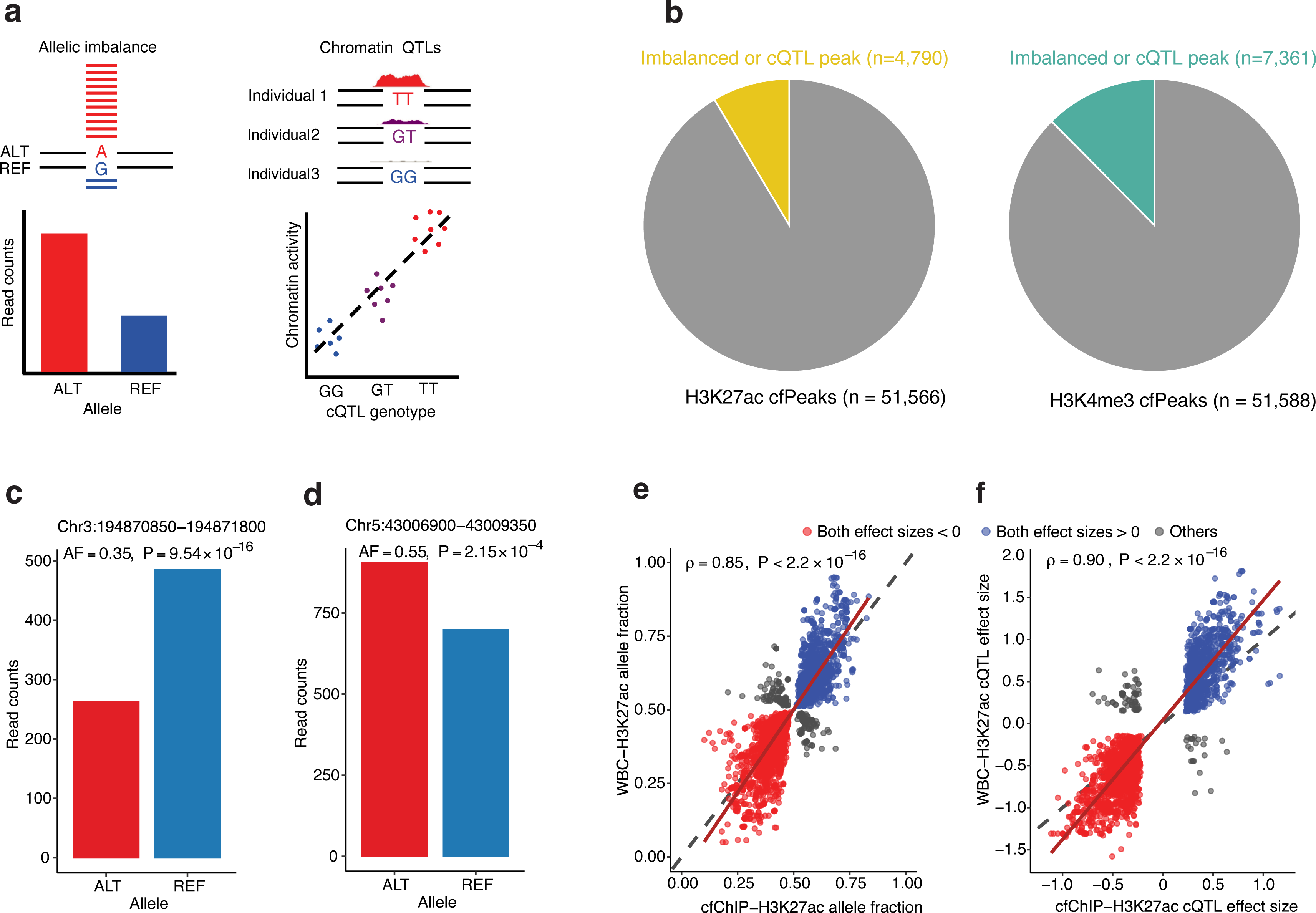
Cell-free chromatin QTL discovery. **(a)** Illustration of allelic imbalance (AI) and chromatin quantitative trait loci (cQTLs). AI in epigenomic data is detected based on differential sequencing read counts between alleles at a heterozygous SNP. cQTLs are SNPs whose genotypes are correlated with chromatin states. **(b**) Proportion of H3K27ac and H3K4me3 peaks with AI or cQTLs identified by a combined test. **(c)** Representative examples of cfChIP-specific allelic imbalance. SNP rs60143196 (Chr3:194870850-194871800) showing differential read counts between reference (REF) and alternative (ALT) alleles aggregated across heterozygous individuals (N=131), with significant reference allele bias (AF = 0.35, P = 9.54 × 10⁻¹⁶). (**d**) SNP rs7729529 (Chr5:43006900-43009350) demonstrating alternative allele bias (AF = 0.55, P = 2.15 × 10⁻⁴) in heterozygous samples (N= 94) with sufficient read coverage (>10 reads. (**e**) Scatterplot comparing allelic fractions for SNPs with significant allelic imbalance in both H3K27ac cfChIP-seq and WBC ChIP-seq datasets (ρ=0.85, p < 2.2×10⁻¹⁶). (**f**) Scatterplot comparing cQTL effect sizes for SNPs with significant chromatin QTLs in both reference panels (ρ=0.90, p < 2.2×10⁻¹⁶).

To validate the robustness of our cfcQTL findings and assess potential technical artifacts, we compared cfcQTLs to WBC H3K27ac ChIP-seq reference panels (“WBC cQTLs”). For SNPs with significant *cis*-genetic effect in both panels, the allelic fraction (AF) of heterozygous SNPs and the effect sizes of cQTLs were highly correlated between the cfChIP and WBC ChIP data (ρ=0.85 for allelic fractions, ρ=0.90 for effect sizes, both p-values < 2.2×10⁻¹⁶; **Fig. 2e, f**). This strong correlation suggests that the cQTLs discovered from circulating chromatin are reproducible in ChIP-seq data and argues against a high rate of false positives due to cancer-associated copy number alterations, which are absent in WBC.

### Cell-free chromatin QTLs reveal context-dependent genetic effects on gene regulation

To evaluate whether cfcQTLs capture tissue- or context-dependent genetic effects on gene regulation, we analyzed the overlap between cfcQTLs and enhancer annotations from reference epigenomes spanning N=98 tissue/cell types. By grouping enhancer regions (EnhA1, EnhA2, EnhG1, EnhG2) from 98 epigenomes into broader tissue categories^49,74^, we quantified the overlap of QTL-containing peaks across these reference tissues. Compared to WBC cQTLs, cfcQTLs showed a higher proportion of overlap in 11 out of 16 tissue categories, including nearly all non-hematologic tissues (**Fig. 3a**). cfcQTLs exhibited the highest percentage of overlap (81.0%) in “other tissues or organs,” a category encompassing active and genic enhancers from liver, pancreatic islets, placenta, lung, ovary, pancreas, spleen, and fetal tissues. In contrast, WBC cQTLs showed higher overlap percentages in blood-related tissues. These results show that cfcQTLs capture regulatory elements influenced by genetic variants in a range of non-hematologic tissues.

**Fig. 3.**
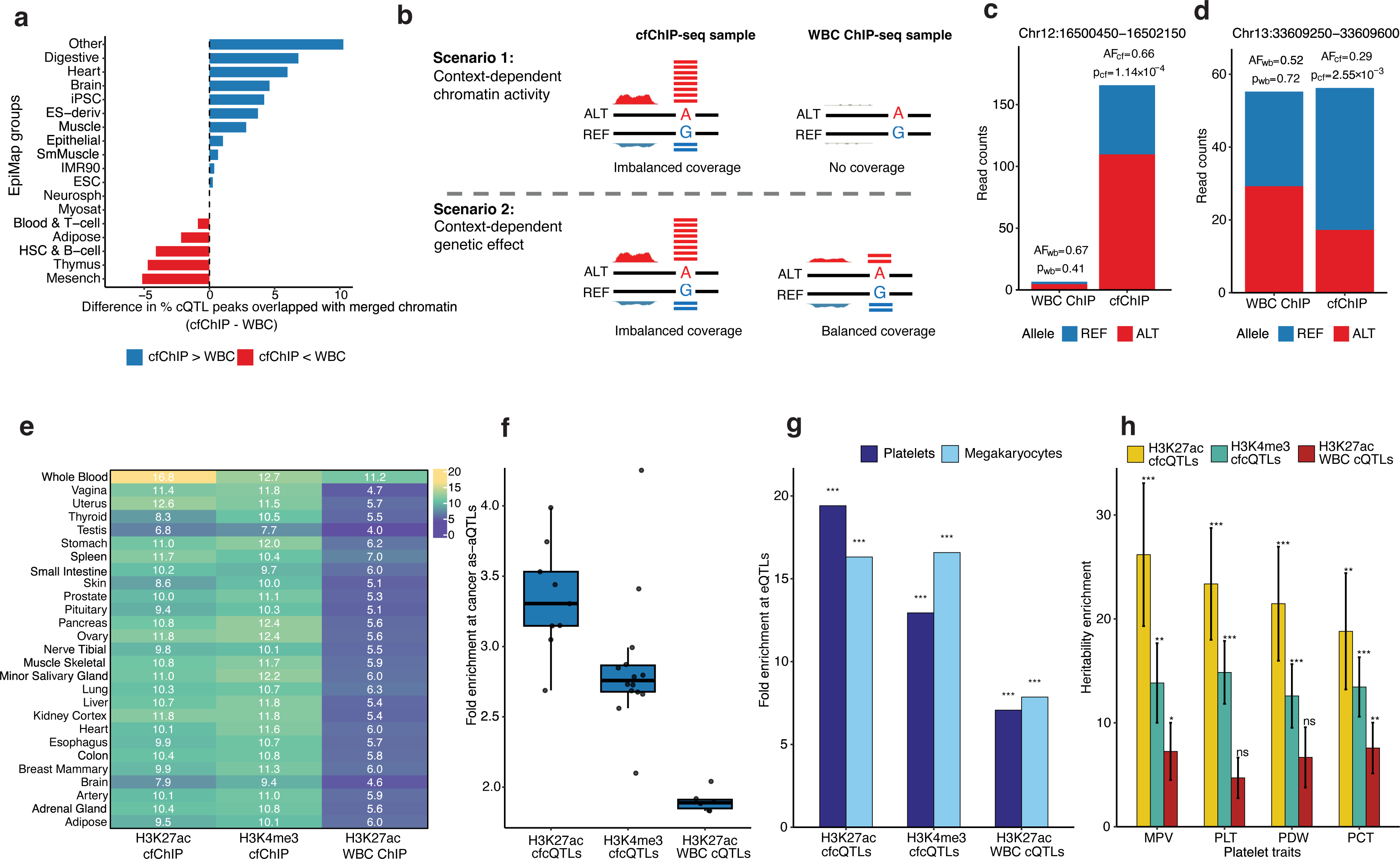
Cell-free cQTLs reveal context-dependent effects on gene regulation. **(a)** Distribution of cQTLs containing peaks within enhancer regions across distinct EpiMap epigenome classifications, showing tissue-specific enrichment patterns. **(b)** Schematic representation of two scenarios where significant cQTLs are detected exclusively in cfChIP-seq. Left panel: cQTLs located in regulatory elements that are not active in white blood cells (WBCs), resulting in absent chromatin peaks in WBC ChIP-seq depicting context-dependent chromatin activity. Right panel: Active chromatin regions detected in both cfChIP-seq and WBC ChIP-seq datasets, with allelic imbalance observed exclusively in cfChIP-seq, indicating context-dependent genetic effects regulation. **(c)** Representative example of cfChIP-specific allelic imbalance at chr12:16500450-16502150, showing minimal coverage in WBC ChIP-seq compared to robust coverage and clear allelic imbalance in cfChIP-seq. **(d)** Example of cfChIP-specific allelic imbalance at chr13:33609250-33609600, showing comparable coverage in both assays but only allelic imbalance in cfChIP-seq. **(e)** Overlap of cfcQTLs and WBC cQTLs with tissue-specific eQTLs in GTEx. Color intensity represents fold enrichment with numerical values indicated within each tile. **(f)** Enrichment for overlap with allele-specific chromatin accessibility sites cancer (as-aQTLs) for H3K27ac cfcQTLs, H3K4me3 cfcQTLs, and WBC cQTLs. Each dot represents one cancer type. **(g)** Enrichment analysis of cfcQTLs in megakaryocyte and platelet eQTLs compared to random genomic background with matched peak characteristics. Statistical significance was established through 10,000 permutations (***P < 2 × 10e-16). **(h)** Stratified LD-score regression analysis identifies significant heritability enrichment of cfcQTLs for platelet-related hematological traits, including mean platelet volume (MPV), platelet count (PLT), platelet distribution width (PDW), and plateletcrit (PCT). Error bars represent standard errors; statistical significance indicated as **P < 0.01, *P < 0.05, and ns (not significant) for P > 0.05.

We considered two models that could explain the detection of cfcQTLs in circulating chromatin but not in WBCs (**Fig. 3b**). The first model—”context-dependent chromatin activity”—involves regulatory elements that are active in cell-free chromatin from non-hematologic tissues but not in white blood cells, such that *cis*-genetic effects can be detected in cfChIP-seq data but not in WBC ChIP-seq data. The second model—”context-dependent genetic effects”—involves regulatory elements that are active in both cfChIP-seq and WBC ChIP-seq (with detectable peaks in both datasets), but where *cis*-genetic variants affect regulatory element activity only in cell-free chromatin context. This scenario could arise from a SNP in a broadly active regulatory element that alters the element’s activity in non-hematologic tissues, but not in WBCs.

Our data revealed examples supporting both models. Supporting the first model, for example, we identified cfcQTLs at peaks with minimal read coverage in WBC ChIP-seq (**Fig. 3c**), consistent with regulatory elements that are active in circulating chromatin but not in WBC. We also identified cfcQTLs exemplifying the second model, where chromatin signals were similar in both reference panels, but genetic effects were only detected in cell-free chromatin (**Fig. 3d**). One such example is rs17403123, a cfcQTL adjacent to *IFRD1*, a gene with comparable expression across multiple tissues, including whole blood, based on GTEx^11^. This variant is an eQTL for *IFRD1* in multiple non-blood tissues but shows no detectable eQTL effect in whole blood (P=0.16 in GTEx). Despite similar chromatin signal intensity at this locus in cfChIP and WBC, allelic imbalance was significant in cfChIP but not in WBC, with a highly significant test for differential allelic imbalance between the two datasets (p=9 x 10^-4^). Additionally, the most significant cfChIP-specific allelically imbalanced regions showed significantly elevated chromatin signal in circulating chromatin compared to WBCs (p=3.3×10⁻¹¹), indicating context-dependent regulatory activity at these elements (**Supplemental Note**).

To quantify the relative frequency of the two models (context-dependent chromatin activity versus context-dependent genetic effects), we analyzed 2,361 peak-SNP associations with allelic imbalance detected exclusively in the cfChIP-seq panel (cfChIP-specific AIs). Applying stratAS^75^ to assess differential allelic fractions between cfChIP and WBC panels, we found that 563 of 2361 (23.8%) peaks exhibited significant differential allelic imbalance (beta-binomial test, FDR < 0.05), consistent with context-dependent genetic regulation. When restricting to regulatory elements with cfChIP-specific AI that are also cfcQTLs, a similar portion (193 of 941, 20.5%) showed differential effects. These observations suggest that a substantial minority of cfcQTLs capture context-dependent genetic effects at regulatory elements that are active in both hematopoietic and non-hematopoietic tissues.

To further determine whether cell-free chromatin QTLs (cfcQTLs) influence transcriptional regulation, we assessed their overlap with previously characterized expression QTLs (eQTLs) from GTEx (v.8)^11^ and allele-specific chromatin accessibility QTLs (as-aQTLs)^24^. cfcQTLs derived from both H3K27ac and H3K4me3 reference panels were significantly enriched for overlap with eQTLs across all 49 GTEx cell lines or tissues. H3K27ac cfcQTLs showed mean enrichment of 9.76-fold (s.d. = 1.83 across tissues), while H3K4me3 cfcQTLs demonstrated 10.62-fold enrichment (s.d. = 1.19), compared to a smaller 5.53-fold enrichment (s.d. = 1.10) for WBC cQTLs (**Fig. 3e, Supplementary Table 4**). Compared to WBC cQTLs, cfcQTLs exhibited substantially higher enrichment for tissue-specific eQTLs (4.62-fold higher for H3K27ac cfcQTLs and 5.12-fold higher for H3K4me3 cfcQTLs; **Fig. 3e, Supplementary Fig. 2**). In addition, cfcQTLs were enriched for sites of allele-specific chromatin accessibility (as-aQTLs) identified across 23 cancer types (**Fig. 3f, Supplementary Table 5).** These findings further indicate that cfcQTLs can capture genetic effects on gene regulation that affect transcription across a variety of tissues, including non-hematologic cell types and cancer tissues.

We investigated whether cfcQTLs reflect transcriptional regulation in normal cell types that contribute to cell-free DNA (cfDNA) outside of the context of cancer. Megakaryocytes, which give rise to platelets, contribute substantially to cfDNA based on cell-type-specific methylation markers^76,77^. Analysis of eQTLs from platelets and induced pluripotent stem cell-derived megakaryocytes (MK) from 185 MK cell lines and 290 blood platelet samples^78^ revealed marked enrichment of cfcQTLs, with H3K27ac showing 19.41-fold and 16.32-fold enrichment for megakaryocytes and platelets, respectively, while H3K4me3 demonstrated 12.94-fold and 16.58-fold enrichment compared to randomly sampled genomic regions. These enrichments exceeded those observed for WBC cQTLs (**Fig. 3g**). Consistent with these findings, stratified linkage disequilibrium score regression (LDSC) demonstrated substantially greater heritability enrichment of cfcQTLs compared to WBC cQTLs for platelet traits including mean platelet volume, platelet count, plateletcrit, and platelet distribution width (**Fig. 3h**). Additionally, cfPeaks not overlapping WBC peaks were enriched for proximity to genes involved in platelet degranulation (**Supplementary Fig. 3**). These findings further support that cfcQTLs capture genetic determinants of gene regulation across multiple cell types that contribute to circulating chromatin.

### Cell-free CWAS identifies novel genetic determinants of cancer risk

To investigate whether cell-free chromatin could identify disease-associated genetic effects on gene regulation, we performed cistrome-wide association studies (CWAS) using our cfChIP reference panels (cfCWAS). This approach identifies genetically determined epigenomic features associated with disease risk by building predictive models of cfChIP-seq signal intensity and allelic imbalance, which are then imputed into GWAS summary statistics^8^. We constructed integrative prediction models for total and allele-specific peak intensity as a function of SNP genotypes within 25kb of a peak. Through fivefold cross-validation, we identified 5,196 of 31,999 (16.23%) H3K27ac cfPeaks and 8,173 of 31,294 (26%) H3K4me3 cfPeaks that showed significant correlation between SNP genotypes and peak intensities at 5% false discovery rate (FDR) in held-out samples.

We first applied cfCWAS to publicly available summary statistics from cancer GWAS of seven common cancers with large case-cohorts ^69,79–85^ and well-powered studies of major non-oncologic diseases, including autoimmune and neurologic diseases such as Crohn’s disease^86^, inflammatory bowel disease (IBD)^87^, rheumatoid arthritis^88^, ulcerative colitis^89^, psoriasis^90^, schizophrenia^91^, and Alzheimer’s disease^92^. We controlled for multiple hypothesis testing using FDR correction across all CWAS models with significant cross-validation performance (CV q < 0.05 per trait), selecting the best-performing model for each peak-trait combination, and identified 150 H3K4me3 and 153 H3K27ac associations for cancer types, and 210 H3K4me3 and 245 H3K27ac associations for non-cancer diseases (**Fig. 4a-b, Extended Fig. 3a-b, Supplementary Table 6**). Applying the conventional genome-wide significance threshold (P < 5×10⁻⁸) to peaks with significant CWAS models yielded 98 H3K4me3 and 88 H3K27ac associations for cancer traits, and 117 H3K4me3 and 124 H3K27ac associations for non-cancer traits. For both histone marks, most of the significant cfCWAS loci (defined as genomic regions within ±500kb of significant cfCWAS associations) overlapped regions containing significant GWAS associations. However, cfCWAS also identified genomic loci with significant associations that did not contain genome-wide significant SNPs in the corresponding GWAS, highlighting the increased statistical power of the CWAS framework to detect novel risk loci **(Fig. 4c-d, Extended Fig. 3c)**.

**Fig. 4.**
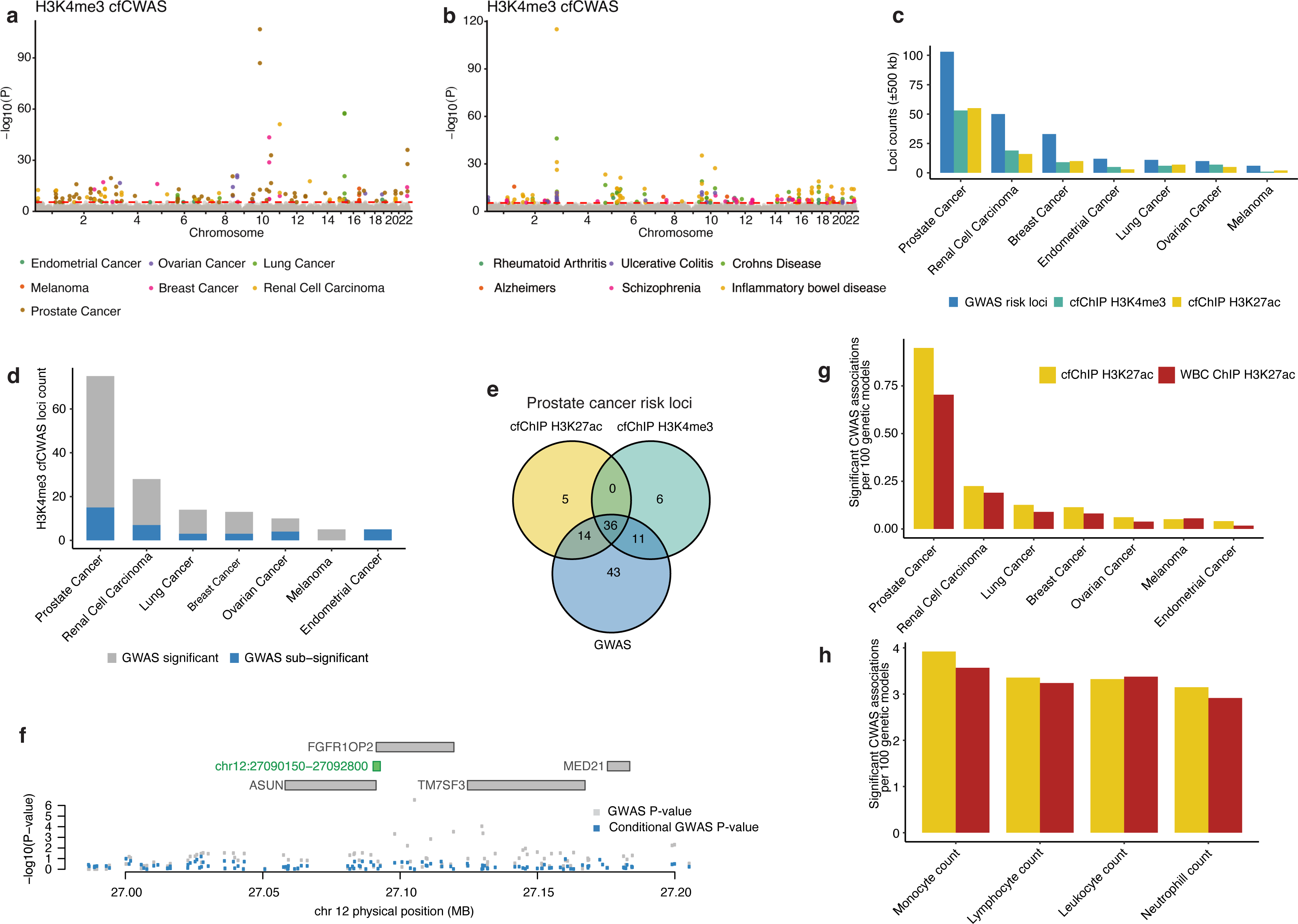
CWAS identifies novel cancer risk loci using cell-free epigenomic reference panels. **(a**) Manhattan plots depicting genome-wide significant associations (−log₁₀(P)) for multiple cancer types using H3K4me3 cfChIP-seq reference panels, with colors denoting different cancer types as indicated in the figure legend. **(b)** Corresponding Manhattan plots for multiple non-cancer diseases based on H3K4me3 cfChIP-seq reference panels. **(c)** Quantitative assessment of significant associations identified across seven cancer types, comparing GWAS (blue) versus CWAS approaches. **(d)** Analysis of loci with significant H3K4me3 cfCWAS associations, distinguishing between GWAS-significant loci (gray) and sub-threshold GWAS loci (blue) that achieved statistical significance only through the enhanced statistical power of the CWAS framework **(e)** Venn diagram illustrating the distinct and overlapping prostate cancer risk loci identified by different molecular QTL approaches (H3K27ac cfChIP-seq, H3K4me3 cfChIP-seq) and conventional GWAS. **(f)** Regional association plot of chromosome 12p11.23 demonstrating attenuation of prostate cancer GWAS signal after conditioning on H3K4me3 cfCWAS signal. **(g)** Quantitative comparison of CWAS performance using cfChIP H3K27ac versus WBC ChIP H3K27ac reference panels across seven cancer types. The y-axis represents the percentage of genetically determined chromatin accessibility regions harboring significant disease associations. **(h)** Corresponding comparisons between cfChIP H3K27ac and WBC ChIP H3K27ac reference panels for four blood-cell-counts GWAS traits.

To validate that novel cfCWAS associations represent true risk loci, we examined whether loci containing these associations achieved genome-wide significance in subsequent larger GWAS. For prostate cancer, we identified 11 cfCWAS loci (H3K4me3 and H3K27ac) lacking genome-wide significant associations in the original GWAS^93^. Five of these 11 loci (45.5%) achieved genome-wide significance in a larger prostate cancer GWAS (**Supplementary Table 7**), with cfcQTLs showing strong linkage disequilibrium (r² = 0.80-0.92) to the lead SNPs identified in subsequent GWAS. H3K27ac and H3K4me3 detected both shared and private cfCWAS associations (**Fig. 4e**, **Supplementary Table 7**) a subset of which were not significant in previous GWAS, including peaks near *FGFR1OP2* (H3K4me3) and *SREBF1* (H3K27ac)—genes linked to pathways implicated in prostate cancer^94–98^ (**Fig. 4f, Extended Fig. 3e)**. Collectively, these observations demonstrate that cfCWAS enhances power for novel risk locus discovery by leveraging genetically determined chromatin features rather than testing all common variants without functional context.

To further evaluate the generalizability of the cfCWAS framework beyond cancer, we expanded our analysis to include 3,900+ GWAS summary statistics from ∼360,000 UK Biobank samples^99^, encompassing a diverse spectrum of human traits and diseases, including cardiovascular, metabolic, neurological, psychiatric, autoimmune, and anthropometric phenotypes. Combined with the curated cancer and disease GWAS described above, H3K4me3 cfCWAS identified significant associations across 1,011 unique traits, involving 4,891 unique cfcQTLs and 4,973 unique peaks (**Supplementary Table 8**). H3K27ac cfCWAS analysis identified associations across 995 unique traits (977 UKBB traits and 18 non-UKBB traits), involving 4,407 unique cfcQTLs and 4,546 unique peaks (**Supplementary Table 9)**. These findings underscore that cfCWAS identifies regulatory associations across diverse disease domains beyond the cancer context of the reference panels.

We next compared how efficiently circulating chromatin CWAS (cfCWAS) captured disease-associated regulatory elements compared to CWAS using WBC H3K27ac profiles (WBC CWAS). We accounted for differences in sample size (N=478 individuals for WBC vs N=303 individuals for cfCWAS), which affects power for discovering associations, by quantifying trait associations per genetically determined peak. cfCWAS consistently yielded a higher proportion of significant peak-trait associations across all seven cancer types. For cancer risk, cfCWAS identified 93 significant associations from 9,802 genetically determined peaks (9.49%), while WBC CWAS detected 204 significant associations from 28,990 peaks (7.04%) (**Fig. 4g**). This pattern persisted across non-cancer diseases, including Inflammatory Bowel Disease (IBD), Schizophrenia, and Crohn’s disease, where cfCWAS consistently yielded higher rates of significant peak-trait associations despite analyzing a smaller set of genetically determined peaks (**Extended Fig. 3d, Supplementary Table 6**). Cochran–Mantel–Haenszel tests confirmed significantly higher association efficiency (measured by associations per genetically determined peak) for cfCWAS than WBC CWAS across cancer traits (p=0.003, OR=1.34) and non-cancer diseases (p=1.8×10⁻⁵, OR=1.40). In contrast, for blood-related traits including monocyte count, neutrophil count, and lymphocyte count, cfCWAS and WBC CWAS showed similar proportions of significant associations per genetically determined peak (**Fig. 4h**). These findings show that cfCWAS provides an efficient approach for identifying functional regulatory elements associated with disease risk, particularly for conditions involving non-hematologic tissues.

### Cell-free chromatin QTLs capture genetic effects on developmental regulatory programs linked to disease risk

A potential advantage of cancer-derived circulating chromatin for studying complex traits and diseases is that cancer reactivates developmental and “stemness” gene regulatory programs that are not observable in differentiated adult tissues^100–103^. We hypothesized that circulating cancer chromatin provides access to regulatory elements from developmental programs that are relevant to genetic risk of disease. To test this hypothesis, we annotated cfPeaks and cfcQTLs based on their overlap with regulatory elements active during distinct developmental stages^49^. cfPeaks were strikingly enriched at non-adult regulatory elements compared to WBC peaks. We observed 21.2-fold enrichment at developmental-specific sites (N=1,170 cfPeaks; p < 10⁻³⁰⁰), 17.2-fold at stem cell-specific sites (N=2,040 cfPeaks; p < 10⁻³⁰⁰), and 5.44-fold at fetal-specific sites (N=139 cfPeaks; p = 2.89×10⁻²⁵). In contrast, adult-specific sites were significantly depleted (0.81-fold; p = 2.95×10⁻²⁵; **Fig. 5a; Extended Fig. 5a; Supplemental Note**).

**Fig. 5.**
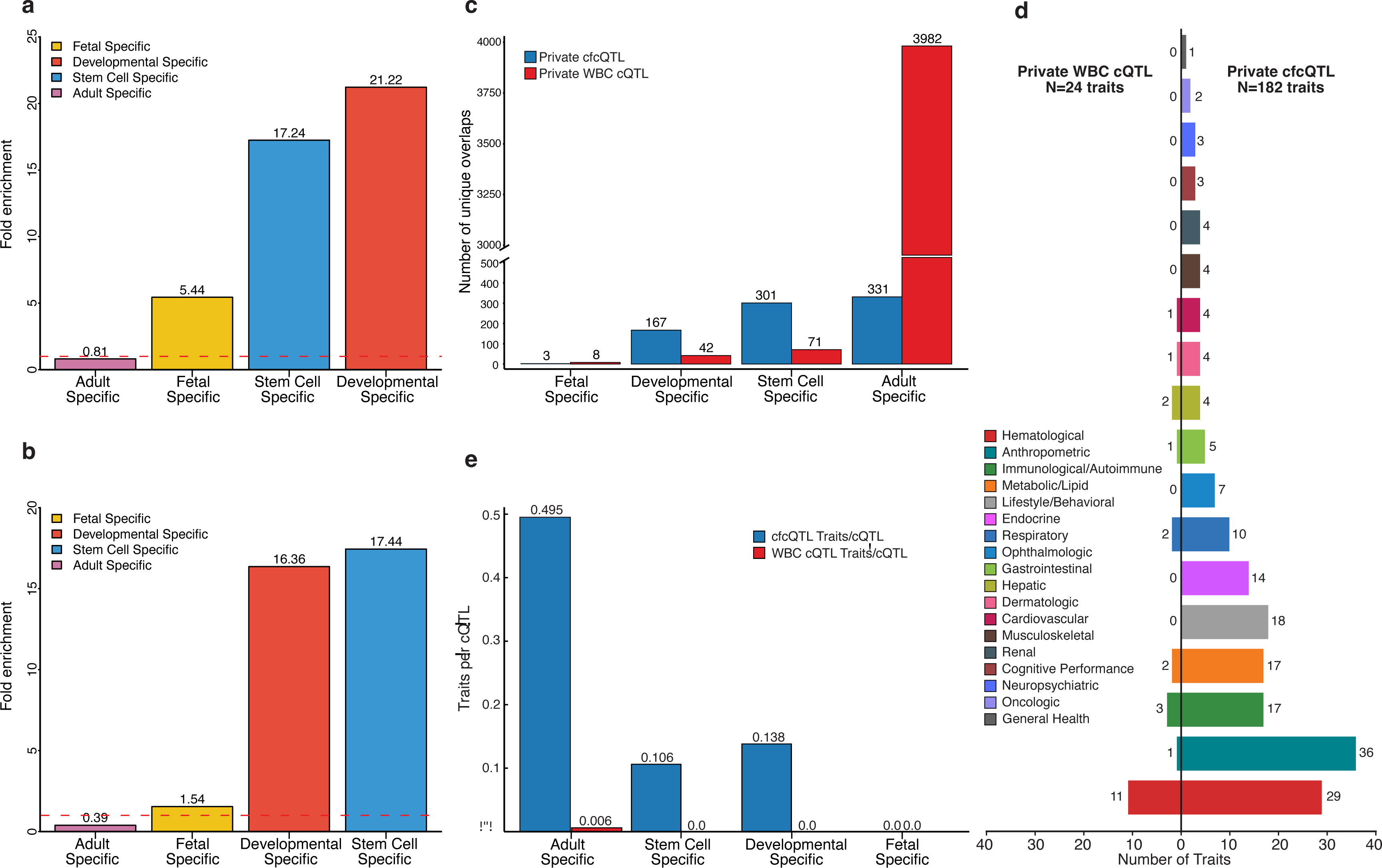
Cell-free chromatin QTLs capture genetic effects on developmental regulatory programs. **(a)** Enrichment of cfPeaks at regulatory elements stratified by developmental stage relative to WBC peaks. **(b)** Enrichment of H3K4me3 cfcQTLs at developmental-restricted regulatory elements relative to WBC cQTLs. **(c)** Distribution of private cfcQTLs (detected exclusively in cfChIP-seq) and private WBC-cQTLs (detected exclusively in WBC ChIP-seq) across developmental categories of regulatory elements. **(d)** Trait category distribution for private cfcQTLs (N=182 traits) versus private WBC cQTLs (N=24 traits) across developmentally stratified regulatory categories. **(e)** Regulatory efficiency of private cfcQTLs versus private WBC-cQTLs across developmental categories, showing traits association per cQTL.

cfcQTLs showed similar enrichment patterns (**Fig. 5b; Extended Fig. 5b)**, with 16.4-fold enrichment at developmental-specific elements (N=167 cfcQTLs; p = 6.68×10⁻⁷⁹), 17.4-fold at stem cell-specific elements (N=301 cfcQTLs; p = 4.51×10⁻¹⁴⁴), and 4.16-fold enrichment at regulatory elements positively correlated with circulating tumor DNA levels^48^ (“CREs”, p = 5.35×10⁻²²¹; **Supplemental Note**). cfcQTLs overlapping CREs were enriched near genes related to cancer and development (**Supplementary Fig. 5**). These findings demonstrate that circulating chromatin captures developmental regulatory elements reactivated in cancer, revealing genetic effects on these otherwise inaccessible regulatory programs.

To evaluate the developmental regulatory landscape captured by cfChIP-seq, we compared “private cfcQTLs” (present only in cfChIP-seq) and “private WBC-cQTLs” (present only in WBC ChIP-seq). Private cfcQTLs showed substantial representation across developmental categories: 3 fetal-specific, 167 developmental-specific, 301 stem cell-specific, and 331 adult-specific. In contrast, private WBC-cQTLs were almost entirely adult-specific (3,982 of 4,103; 97%), with minimal representation of developmental regulatory elements (**Fig. 5c; Supplementary Fig. 6-7**). We examined disease associations by overlapping these developmentally stratified private cQTLs with our CWAS results from UKBB phenotypes. Private cfcQTLs in developmentally stratified regulatory regions were associated with diverse traits (N=182 total) spanning oncological and non-oncological phenotypes, including neurological, endocrine, cardiovascular, hepatic, renal, and lipid metabolism traits (**Fig. 5d; Extended Fig. 5c; Supplementary Table 10-11**). In contrast, private WBC-cQTLs demonstrated fewer (N=24) and more restricted disease associations, predominantly linking to hematological and immune system traits (**Fig. 5d; Extended Fig. 5d; Supplementary Table 12-13; Supplemental Note**). Developmental-specific elements generally showed greater trait specificity (less pleiotropy) for private cfcQTL-associated traits than compared to adult-specific elements (**Supplementary Fig. 9)**.

To assess trait associations per cQTL, we compared private cfcQTLs and private WBC-cQTLs for their ability to associate with traits across developmentally stratified regulatory categories. Overall, across these categories combined, private cfcQTLs demonstrated substantially higher regulatory efficiency (16.6% overall rate; 133 peak-trait associations from 802 cfcQTLs) compared to private WBC-cQTLs (0.2% rate; 8 associations from 4,103 cQTLs), representing an 83-fold difference. Notably, cfcQTLs were associated with more unique phenotypes even in adult-specific regions, associating with 164 unique traits compared to 24 for WBC-cQTLs—a 6.6-fold difference (**Supplementary Table 12,14)**. Overall, across developmentally stratified regulatory regions, private cfcQTLs associated with 182 unique traits compared to 24 for private WBC-cQTLs (**Fig. 5d, Supplementary Table 10,12**), yielding 39-fold more traits per variant on average (0.2269 vs 0.0058 traits per cQTL; **Fig. 5e; Supplementary Fig. 10**).

Collectively, these findings demonstrate that cfChIP-seq captures genetic effects on regulatory elements across diverse developmental stages, tissues, and disease states that are largely inaccessible through conventional WBC ChIP-seq epigenomic profiling.

### cfChIP captures the regulatory effects of *TERT* promoter mutation

Having established that circulating chromatin can reveal the effects on regulatory activity of germline variants, we asked whether it could assess the impact of non-coding somatic mutations in cancer, which are common but difficult to characterize functionally^104–107^. We considered the *TERT* promoter region, which harbors recurrent mutations that create a novel transcription factor binding site recognized by ETS family transcription factors, increasing telomerase expression in multiple cancer types^108–110^. We examined H3K4me3 cfChIP-seq data from plasma samples of cancer patients in our cohort, focusing on the canonical chr5:1,295,228 C>T (C228T)^108^ C to T mutation within the *TERT* promoter (**Fig. 6a**). From cfChIP-seq reads, we identified examples of *TERT* promoter mutations that were absent in healthy plasma samples. Analysis of the variant allele fraction (VAF) revealed H3K4me3 cfChIP-seq fragments were disproportionately enriched for the *TERT* promoter mutation relative to the estimated circulating tumor DNA (ctDNA) content in each sample. For example, in a melanoma sample, we observed a VAF of 80% despite a ctDNA fraction of only 19%; **Fig. 6b**). The disproportionately high enrichment of the mutant allele in H3K4me3 cfChIP-seq data compared to the low portion of tumor-derived DNA indicates preferential enrichment of histone modification at the mutated promoter. This finding is consistent with the observation that the mutant allele increases promoter activity. This proof-of-concept analysis demonstrates that cfChIP-seq can detect cancer-associated non-coding mutations and capture their functional consequences on regulatory element activity directly from patient plasma samples.

**Fig. 6.**
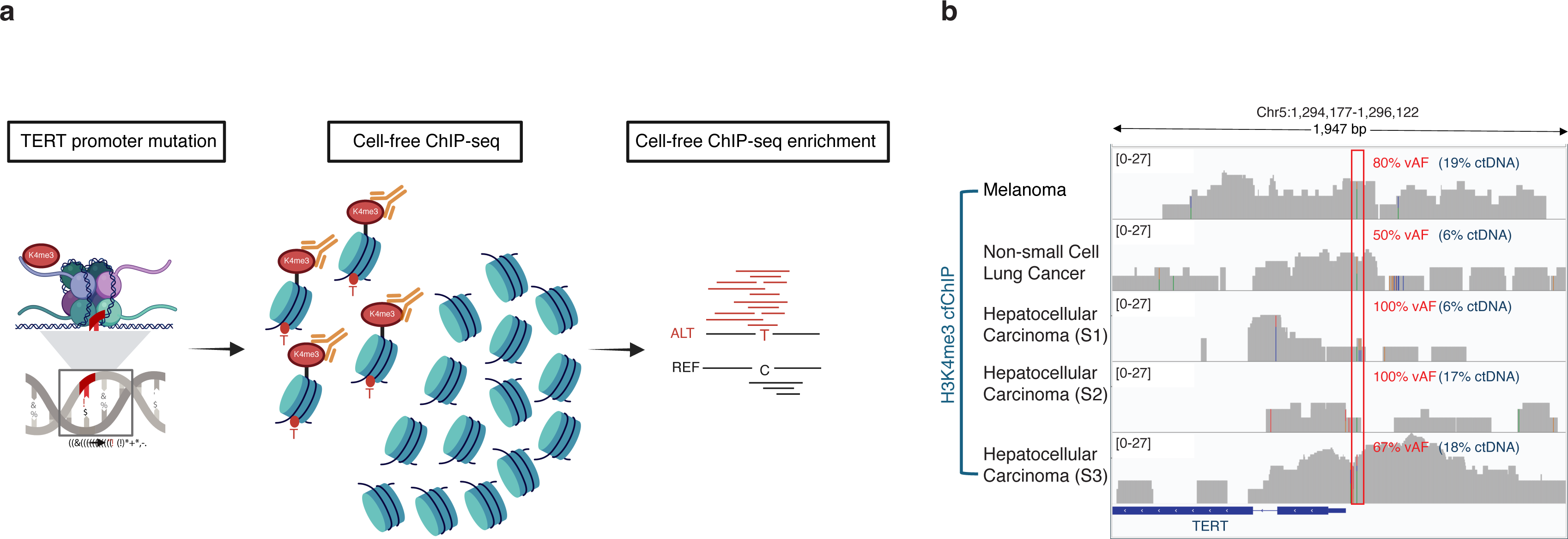
Cell-free ChIP-seq enriches activating *TERT* promoter mutations. **(a)** Schematic illustration of *TERT* promoter mutation detection through cfChIP-seq. The chr5:1,295,228 (C228T) C>T mutation creates a novel ETS transcription factor binding site, enhancing promoter activity, reflected by increased levels of H3K4me3. Schematic figure created with Biorender. **(b)** H3K4me3 cfChIP-seq signal profiles at the *TERT* promoter locus (Chr5:1,294,177-1,296,122) in plasma samples from patients with melanoma, non-small cell lung cancer (NSCLC), and hepatocellular carcinoma (three independent samples, S1-S3). For each cancer type, the variant allele fraction (VAF) in cfChIP-seq reads substantially exceeds the estimated circulating tumor DNA (ctDNA) fraction, demonstrating H3K4me3 enrichment at the mutant allele.

## Discussion

In this study, we demonstrate that circulating chromatin is a powerful substrate for measuring the effects of genetic variants on gene regulation and linking these effects to disease risk. To our knowledge, this is the first study to leverage liquid biopsy to discover chromatin QTLs. This approach has major advantages for studying genetically complex human traits and diseases. Foremost, it can sample chromatin, including developmentally restricted regulatory elements, from non-hematologic tissues from peripheral blood, revealing context-dependent effects of variants on gene regulation that are not present in WBCs. Our comparison of circulating chromatin and WBCs indicates that such context-dependent effects are common, consistent with emerging literature^7–9,18^. In addition, cfcQTL discovery can be performed on a large scale, given the ease of obtaining blood plasma. This will allow major upscaling of cQTL discovery efforts in non-hematologic tissues, which have so far been limited in number and size due to the need to obtain surgical, biopsy, or post-mortem tissues from many individuals.

A key novelty of our approach is that cfChIP-seq enables the study of genetic determinants of gene regulation across tissues and developmental stages that are difficult to access but relevant to disease. We found that cfChIP-seq-derived peaks capture regulatory elements across diverse tissues beyond those detected in white blood cell ChIP-seq data. Both cfPeaks and cfcQTLs demonstrated striking enrichment at non-adult regulatory elements, with developmental-specific sites showing 21.2-fold and 16.3-fold enrichment, respectively, stem cell-specific sites showing 17.2-fold and 17.4-fold enrichment, while adult-specific sites were significantly depleted. Critically, by sampling chromatin from cancer patients, we captured regulatory elements that are not active in adult differentiated tissues but are reactivated in cancer. This advantage enabled us to identify developmentally restricted regulatory elements associated with a wide range of traits and diseases, including anthropometric, behavioral, metabolic, and cardiovascular phenotypes. This capability is particularly advantageous given that a substantial fraction of disease-associated GWAS loci lack explanatory eQTLs in commonly profiled tissues^11,17,111^, often because the relevant regulatory activity occurs only in specific developmental stages or disease states that are difficult to access through conventional approaches.

cfcQTLs were enriched for known eQTLs and chromatin accessibility QTLs from non-blood tissues, confirming that our approach captures genetic effects on gene regulation and expression across a range of tissues. Notably, ∼20% of cfChIP-specific allelic imbalance exhibited statistically significant differential allelic fractions relative to WBCs, providing evidence for context-dependent genetic effects where the same regulatory element is subject to different genetic influences in hematopoietic versus non-hematopoietic contexts. This finding demonstrates that context-dependent genetic effects on gene regulation, which are a key feature of genetically complex diseases, are detectable in circulating chromatin.

We applied the cistrome-wide association study framework (cfCWAS) and demonstrated that it effectively captures genetic effects on chromatin that mediate disease risk. Through systematic analysis of 3,900+ GWAS summary statistics, we identified significant associations across 1,011 unique traits spanning cardiovascular, metabolic, neurological, psychiatric, autoimmune, and anthropometric phenotypes. cfCWAS detected sub-threshold GWAS associations that validated in larger subsequent GWAS and demonstrated higher discovery efficiency for non-blood phenotypes than WBC CWAS. Notably, developmentally restricted private cfcQTLs demonstrated 39-fold more trait associations per variant on average compared to private WBC cQTLs (22.7 versus 0.58 traits per 100 QTLs) and were associated with182 unique phenotypes compared to 24 for WBC-restricted cQTLs. This broad phenotypic coverage demonstrates that circulating chromatin from patients with cancer captures genetic effects on regulatory elements with relevance to non-oncologic diseases and traits.

Finally, we provide a proof of concept for adapting our approach to detect the effects of somatic mutations on regulatory element activity. This capability could provide an *in vivo* functional assessment of how patient-specific mutations in noncoding DNA affect gene regulation in cancer.

Our study has several limitations. Our cfChIP-seq reference panel’s modest sample size constrains statistical power for detecting genetic effects. While cfDNA captures chromatin features from multiple tissues, quantifying the relative contribution of each tissue remains challenging, though emerging deconvolution approaches offer potential solutions^112,113^. Future work will leverage the scalability of cfChIP-seq to enable large-scale studies across diverse patient populations to address the paucity of non-European epigenomic profiling data. The comprehensive cfCWAS resource generated here, linking genetic variants to chromatin activity across thousands of traits, provides a reference for functional interpretation of GWAS findings.

## Data availability

Data sets generated from this study, including cell-free cQTLs and cell-free CWAS results are available at Zenodo: https://doi.org/10.5281/zenodo.17538570. Chromatin states and histone marks related functional annotation/data were obtained from Roadmap Epigenomics web portal https://egg2.wustl.edu/roadmap/web_portal/chr_state_learning.html and Epimap https://compbio.mit.edu/epimap/. UK Biobank summary stats were obtained from: https://www.nealelab.is/uk-biobank. Processed cfChIP-seq data, including BED files containing genomic alignments and peak calls, used in this study are publicly available in the Gene Expression Omnibus (GEO) under accession <u>GSE243474</u>. Due to privacy protections, raw sequencing files are available from the corresponding author upon reasonable request subject to a data use agreement.

## Code availability

Code used for the analyses presented in this paper is available on GitHub: https://github.com/chhetribsurya/cell-free-cQTL-cfCWAS

## Acknowledgments

This work is supported by the US Department of Defense (DoD) (awards W81XWH-21-1-0358 and W81XWH-21-1-0299 to SCB), the National Cancer Institute (U01CA296432 award to SCB), a Prostate Cancer Research Foundation Young Investigator Award (SCB), The Damon-Runyon Cancer Research Foundation Clinical Investigator Award (SCB), and The Fund for Innovation in Cancer Informatics. We are grateful for the generous support of Debbie and Bob First and Mark Donovan. The content is solely the responsibility of the authors and does not necessarily represent the official views of the National Institutes of Health.

## Methods

### Data and quality control

cfChIP-seq data were collected from a prior publication^48^, which included cancer patients treated at the Dana-Farber Cancer Institute (DFCI), the National Cancer Institute (NCI), and Massachusetts General Hospital (MGH), and healthy individuals from the Mass General Brigham Biobank. ChIP-seq peak location information and genomic alignments of all sequenced fragments can be retrieved through GEO under accession number GSE243474. Request for raw sequencing data should contact the corresponding authors of the publication.

Sequence data were processed as described previously^48^ . In brief, sequencing reads were aligned to the hg19 human genome build using Burrows-Wheeler Aligner version 0.7.1740, followed by discarding non-uniquely mapping and duplicated reads. ChIP-peak calling was conducted using MACS version 2.1.1.2014061641 with a threshold of false discovery rate (FDR) < 0.01. BED files containing fragment location information were generated using BEDTools version 2.29.2 with the -bedpe flag set. An on-target to off-target enrichment ratio for H3K27ac and H3K4me3 was calculated to assess immunoprecipitation specificity^48^, and we included cfChIP-seq samples with the product of the number of unique fragments and the enrichment ratio > 2 x 10^6^ for all analyses conducted in this study.

Sets of consensus peaks for H3K27ac and H3K4me3 cfChIP-seq were constructed by dividing the whole genome into 50-bp windows and keeping any windows with peaks in > 5 of the samples. Then, retained windows were buffered by 100 bp and merged to construct sets of 51,542 H3K27ac peaks and 53,627 H3K4me3 peaks.

### Genotype imputation

Genotypes at 5,495,776 autosomal SNPs with minor allele frequency > 5% in the Haplotype Reference Consortium (HRC, v1.191)^114^ were imputed. We first merged BAM files from epigenomic datasets for each individual using SAMtools merge then run through STITCH version 1.6.2^115^ with the following parameters: *k* = 10, ngen = 1,240, niterations = 40, method = diploid (https://hub.docker.com/r/stefangroha/stitch_gcs/tags). The imputation reference panel was retrieved from phase 3 of the 1000 Genomes Project, containing haplotypes information of 2,505 individuals^116^.

Quality assessment for imputed genotypes from cfChIP-seq samples was conducted as detailed in Ref^48^. To validate BAM files from each individual were correctly assigned, we called genotypes at 100,000 SNPs using bcftools mpileup and call functions (v1.9) and tested the pairwise correlation using bcftools gtcheck function across all files. No samples were based on the clustering results.

### EpiMap reference epigenome curation and comparison with cell-free chromatin

BED files for 98 reference epigenomes (EIDs), annotated with chromatin states from the 18-state Roadmap model, were obtained from the EpiMap repository^49,74^ (https://egg2.wustl.edu/roadmap/data/byFileType/chromhmmSegmentations/ChmmModels/core_K27ac/jointModel/final/all.dense.browserFiles.tgz). H3K27ac-enriched regions were defined for each EID by merging the chromatin states EnhA1, EnhA2, TssA, EnhG1, and EnhG2 using the bedtools merge function. Similarly, H3K4me3-enriched regions were defined by merging states corresponding to TssA, TssFlnk, TssFlnkU, TssFlnkD, and TssBiv. Epigenomes were further categorized based on whether they represented blood-derived or non-blood tissues.

To quantify the overlap between cell-free chromatin and reference chromatin states, we used bedtools intersect to calculate the number of histone mark-enriched regions that overlapped with the consensus peak set of the corresponding assay. Overlap percentages were calculated by dividing the number of overlapping regions by the total number of regions across all EIDs.

To assess tissue specificity, we defined a tissue-specificity score for each consensus peak as the proportion of EIDs in which the peak was present. Peaks with scores in the bottom 25th percentile were denoted as tissue-specific peaks. For each EID, we then counted the number of tissue-specific peaks overlapping its histone mark-enriched regions. These counts were grouped by blood versus non-blood tissue categories to assess differences across cfChIP-seq and WBC ChIP-seq assays.

Statistical differences in overlap between assays were evaluated using the Wilcoxon rank-sum test. To compare enrichment patterns between blood and non-blood EIDs, we calculated the odds ratio of tissue-specific peaks overlapping histone mark-enriched regions in one category versus the other for cfChIP-seq and WBC ChIP-seq assays.

### Genomic browser views of cfChIP-seq data

Normalized cfChIP-seq read counts at selected cancer diagnostic genes and GWAS risk loci were visualized with Gviz version 1.50.0 on R version 4.4.3. We selected three cfChIP-seq samples with the highest ctDNA level for each cancer type among prostate cancer, breast cancer, colorectal cancer, and neuroendocrine prostate cancer. Three WBC ChIP-seq samples were randomly selected to match the sample size. Data from samples representing the same cancer types or WBC were aggregated and overlaid on a single track using the OverlayTrack function.

### Genetic modeling of cell-free chromatin features

Peak intensity of cell-free chromatin for H3K4me3 and H3K27ac were modeled based on cis-SNP genotypes by a Snakemake pipeline available at https://github.com/scbaca/cwas. Allelic imbalance analysis and cis cQTL mapping were conducted for each consensus peak across both histone modifications.

### Allelic imbalance analysis for allele-specific peak intensity

The Snakemake pipeline adopts stratAS^24,75^ to identify imbalance of ChIP-seq reads on heterozygous SNP alleles. To increase power for unbiased imbalance testing, the pipeline incorporated the following steps: 1) imputed SNP genotype were phased with Eagle2^117^ by using Sanger Imputation Service (https://imputation.sanger.ac.uk/) to count reads on nearby heterozygous alleles on the same haplotype; 2) BAM files were processed by the WASP pipeline^118^ to correct for biases in allele-specific mapping; 3) Processed BAM files were later prepared using ASEReadCounter from the Genome Analysis Toolkit (v3.8103)^119^; 4) copy number profiles for each individual were estimated from off-target reads with CopywriteR^120^ to infer the overdispersion parameter 𝜌, which is a locally defined, per-individual sequence read correlation parameter to adjust for overdispersion in regions of copy number alterations required by stratAS.

stratAS models read counts from heterozygous SNPs by assuming a beta-binomial distribution. Given a tested chromatin peak, read counts for an individual 𝑖 overlapping a heterozygous site 𝑗 were 𝑅_alt, 𝑖_|𝑅_ref, 𝑖_ BetaBin(𝜋_𝑗_, 𝜌_𝑖𝑗_), where 𝜋 is the mean allelic ratio. Only SNPs within the corresponding peak were included for the imbalance test in this analysis. False discovery rates (FDR) of allelic imbalance P values were calculated with the qvalue R package (v2.18) to adjust for multiple testing correction. Significant cfPeaks were regarded as peaks containing one or more SNPs with 𝑞 < 0.05. Default parameters^8^ for estimating the overdispersion parameter 𝜌 and conducting the allelic imbalance test were adopted.

### cQTL mapping

cQTL detection was conducted with QTLtools (v1.2)^66^. Reads per kilobase per million mapped reads (RPKM) values for each consensus peak were calculated using QTLtools quan from each BAM file. A covariate matrix was constructed using QTLtools pca--scale--center. Nominal 𝑃 values of cfPeak-cQTL pair within 25kb of the corresponding peak were calculated using QTLtools cis. FDR correction was used to adjust 𝑃 values (𝑞 < 0.05) for downstream analyses. Default parameters from the pipeline^8^ for RPKM value calculation, covariate matrix construction, and QTL detection were adopted in this analysis.

A combined test to evaluate the combined significance of SNP-peak association, considering both allelic imbalance and cQTLs, was constructed using Stouffer’s method^121^. SNP-peak pairs tested in both analyses were included in the combined test.

### CWAS model construction and peak-trait association testing

Predictive models for chromatin peak intensity based on *cis*-SNP genotypes were constructed with the ‘predict’ flag in stratAS. Total histone modification signal, allelic imbalance, and a combination of the two were modeled using LASSO penalized regression based on genotypes from 1) all variants or 2) the top SNP within 25kb of the corresponding peak center. This corresponds to a total of six model types, including “lasso”, “lasso.as”, “lasso.combined”, “top1.as”, “top1”, “top1.combined”. Pearson correlation between predicted values and true chromatin activity across fivefold cross-validation was used to evaluate prediction accuracy. Significant models were defined by first selecting the model type with the most significant cross-validation *P* value within each peak and then retaining models with cross-validation significance at an FDR of 0.05 across all peaks.

The genetic associations between predicted chromatin activity and GWAS risk were calculated using FUSION^122^. CWAS *Z*-scores were considered as a linear combination between a weight matrix **W** and a vector of SNP-trait association *Z*-scores 𝚭_SNP→trait_. The weight matrix **W** was estimated based on the peak-SNP covariance matrix from the above-described models and the SNP-SNP covariance matrix reflecting linkage disequilibrium. Two-sided *P* values were inferred from the standard normal distribution based on the corresponding CWAS *Z*-scores. Detailed description of mathematical derivation of CWAS Z-scores can be found in the original publication^8^. Significant CWAS associations were defined as peak-trait associations with 𝑃 < 0.05 after Bonferroni correction for all significant predictive models.

### Enrichment analysis for consensus cfPeaks and cfcPeaks

We tested the enrichment of molecular QTL SNPs (e.g., expression QTLs or allele-specific accessibility QTLs) in cfcPeaks using permutation analysis. For each type of molecular QTL, we counted the number of QTL SNPs overlapping cfcPeaks and divided by the total base-pairs covered by cfcPeaks. We then repeated this process on 5,000 equally sized samples of the complete set of consensus cfPeaks to create a null distribution. The ratio of peaks containing QTL SNPs in the observed versus simulated data was used to calculate enrichment and a one-sided P value. Additionally, we compared enrichment to a random background by repeating the same process using random intervals matched to cfcPeaks in terms of size, number, and chromosome.

The enrichment analyses of Gene Ontology terms in consensus cfPeaks and cfcPeaks were conducted using the Genomic Regions Enrichment of Annotations Tool (GREAT)^123^ version 4.0.4 (http://great.stanford.edu/public/html/).

### Regulatory region annotation across developmental stages

Regulatory region annotations were derived from chromatin state segmentation data provided by the Roadmap Epigenomics Project^49^, utilizing 18-state ChromHMM models across 127 reference epigenomes. Analysis focused on histone modification profiles characterizing active regulatory elements, including active transcription start sites, genic and active enhancers, weak enhancers, and bivalent regulatory elements. Following standard ChromHMM nomenclature, the following active regulatory states were extracted: 1_TssA (Active TSS), 2_TssFlnk (Flanking TSS), 3_TssFlnkU (Flanking TSS Upstream), 4_TssFlnkD (Flanking TSS Downstream), 7_EnhG1 (Genic Enhancer 1), 8_EnhG2 (Genic Enhancer 2), 9_EnhA1 (Active Enhancer 1), 10_EnhA2 (Active Enhancer 2), 11_EnhWk (Weak Enhancer), 14_TssBiv (Bivalent TSS), and 15_EnhBiv (Bivalent Enhancer).

The 127 reference epigenomes were stratified into four developmental categories based on tissue origin and differentiation state. From each developmental category, genomic regions exhibiting the active regulatory states described above were extracted, resulting in four distinct regulatory region sets. Developmental-specific regions were defined as the union of fetal and stem cell regulatory regions, with any regions overlapping adult-specific regulatory annotations explicitly excluded to ensure developmental specificity. Regulatory regions exhibiting tissue-specific activity patterns were identified through set-based operations on the fetal, stem cell, and adult regulatory region annotations, establishing four mutually exclusive specificity categories: (1) fetal-specific regions, active exclusively in fetal tissues; (2) stem cell-specific regions, active exclusively in stem cell epigenomes; (3) developmental-specific regions, active in fetal and/or stem cell tissues but absent in adult tissues; and (4) adult-specific regions, active exclusively in adult tissues. BEDTools intersection operations were employed to implement these classifications, ensuring complete mutual exclusivity between categories while providing comprehensive coverage of the regulatory landscape.

### Statistical enrichment analysis and significance testing of cQTLs in regulatory regions

Enrichment of cQTLs from circulating chromatin and WBC derived chromatin within each developmental specificity category was assessed through overlap analysis using BEDTools. The number of cQTLs overlapping regulatory regions in each category was calculated, and statistical significance was evaluated using Fisher’s exact test comparing overlap frequencies between the cQTL set and a size-matched random background. To generate the random background, random sampling was performed from the full set of cQTLs at equivalent significance thresholds, maintaining the same total number of regions as the test set. By matching both sample size and statistical thresholds, this design ensured that any observed enrichment reflected genuine biological specificity rather than artifacts arising from differences in statistical power or genomic distribution.

Fisher’s exact test was implemented using a 2×2 contingency table comparing overlapping and non-overlapping regions between the cQTL set and the random background. Odds ratios along with 95% confidence intervals were computed using the log transformation method, where the standard error of the log odds ratio was calculated as the square root of the sum of the reciprocals of all cell counts in the contingency table: SE (log OR) = √(1/a + 1/b + 1/c + 1/d), where a, b, c, and d represent the four cells of the 2×2 table. The confidence intervals were then obtained by exponentiating the log odds ratio ± 1.96 × SE (log OR). And fold enrichment was computed as the ratio of overlap frequencies. To provide complementary evidence for the functional relevance of identified cQTLs, enrichment within regulatory elements positively correlated with circulating tumor DNA levels^48^ (CREs) was assessed, employing the same statistical framework as the developmental specificity analysis.

### Permutation testing and computational implementation

Permutation testing was performed to validate the robustness of enrichment findings. For each developmental specificity category, 100 independent random samples were generated from the full background set, with each sample matched in size to the test cQTL set. A fixed random seed 50 was used to ensure reproducibility across all analyses. Within each permutation, overlap with regulatory regions was calculated and the corresponding fold enrichment and odds ratio were computed.

All genomic coordinate operations were performed using BEDTools^124^ (v2.31.1) and pybedtools^125^ (v 0.12.0) with the human reference genome assembly GRCh37/hg19. Chromosome coordinates were standardized to include the “chr” prefix, and all genomic intervals were converted to 0-based, half-open coordinate format as required by BED format specifications. Statistical analyses were conducted in Python (v 3.10.18) using scipy.stats (with scipy v1.15.2) for Fisher’s exact tests and NumPy (v2.2.6) for numerical computations. Permutation sampling was implemented using Python’s random module with fixed seeds to ensure reproducibility. Data visualization was performed using Matplotlib (v3.10.6) and R (v4.4.0) (ggplot2 package, v 3.5.2).

---

**Supplementary Fig. 1.**
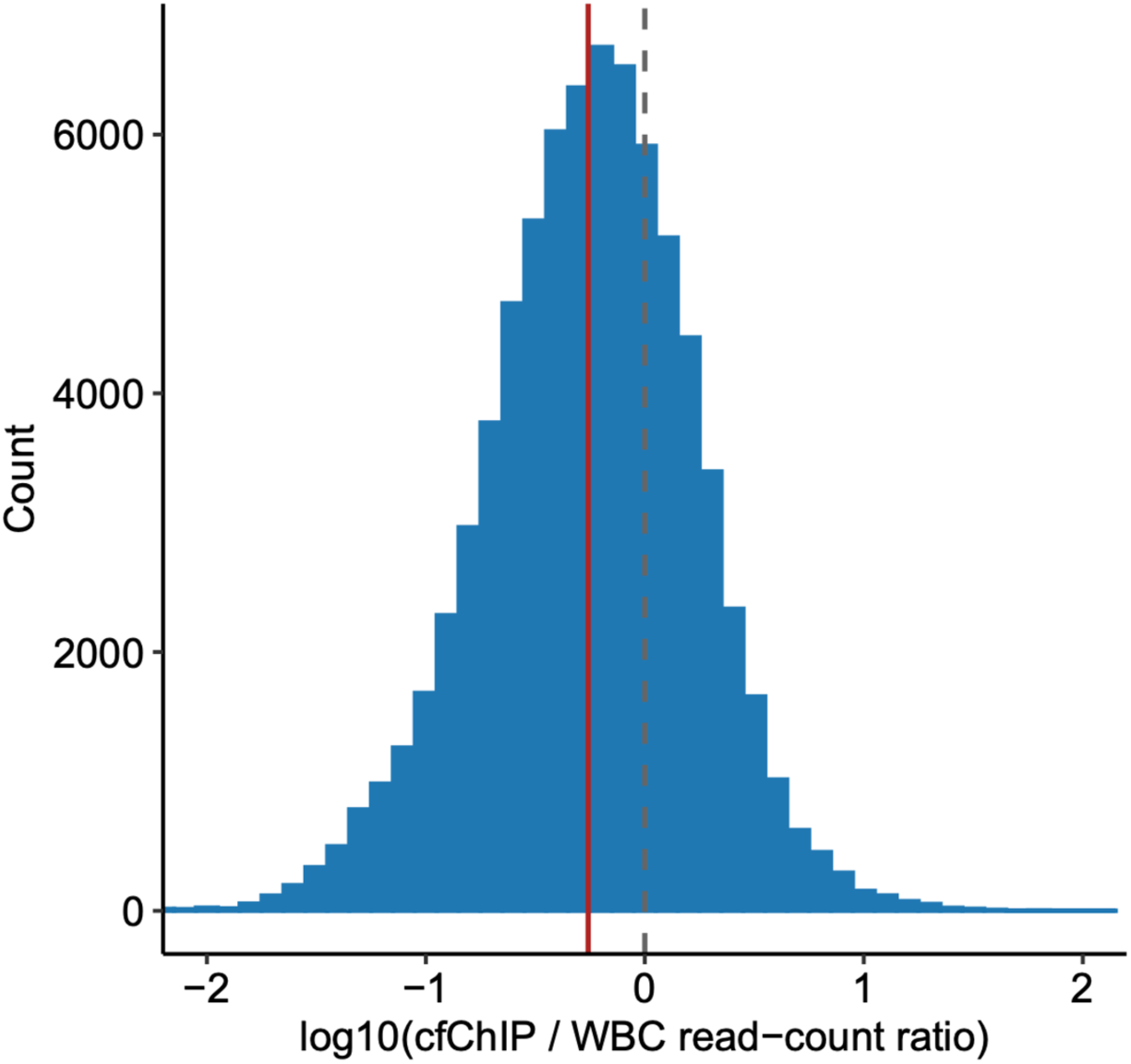
Distribution of cfChIP to WBC read count ratios across all tested peaks. Histogram showing log10-transformed ratios of read counts between cfChIP-seq and WBC ChIP-seq samples across all tested peaks. Red vertical line indicates the mean ratio. Gray dashed vertical line indicates log10 ratio of 0 (equal signal between cfChIP and WBC).

**Supplementary Fig. 2.**
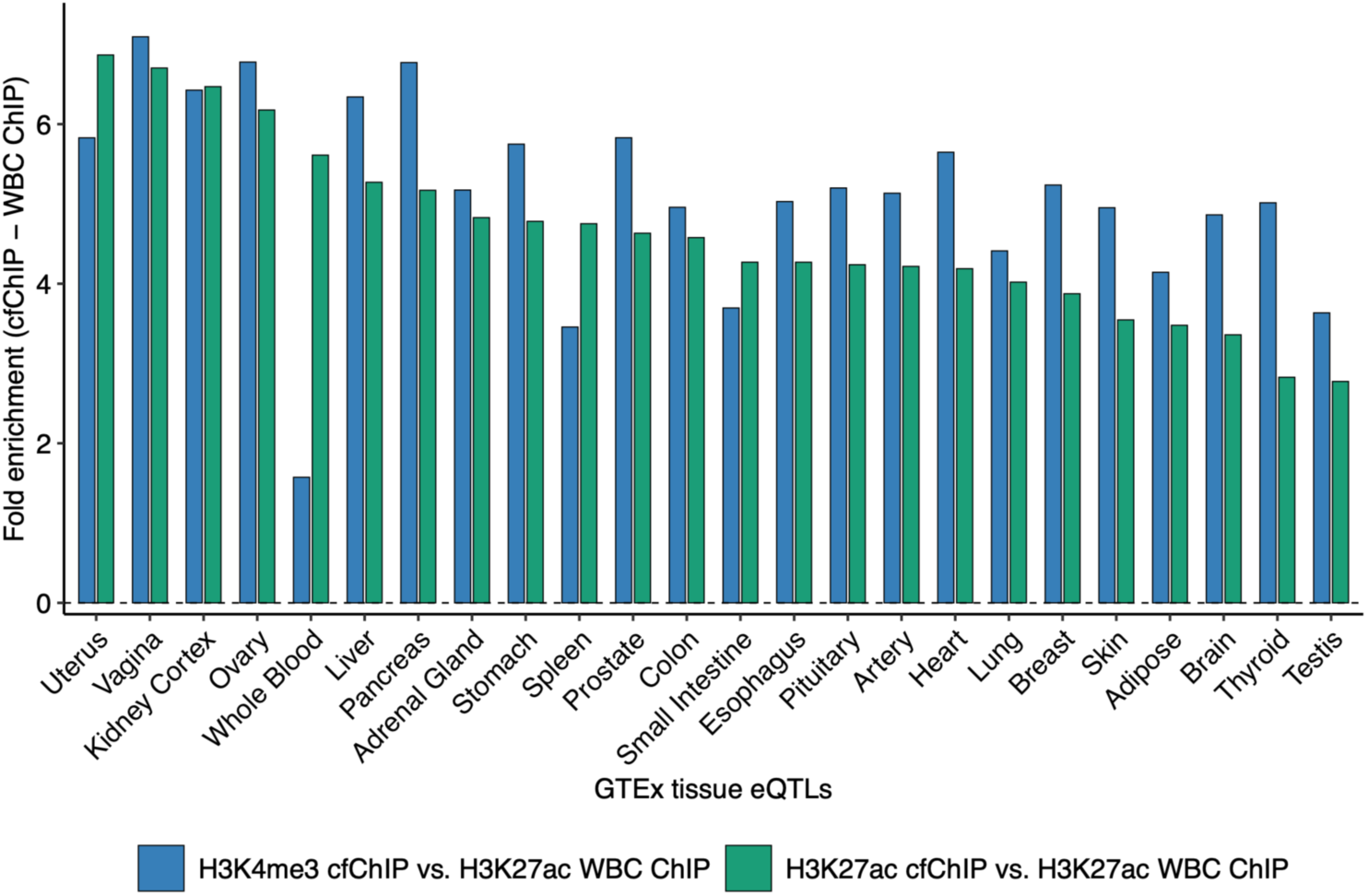
Differential enrichment of GTEx eQTLs between cfChIP cQTLs and WBC cQTLs across tissues. Bar plots showing the differences in GTEx eQTLs fold enrichment of cfChIP cQTLs than of WBC ChIP cQTLs.

**Supplementary Fig. 3.**
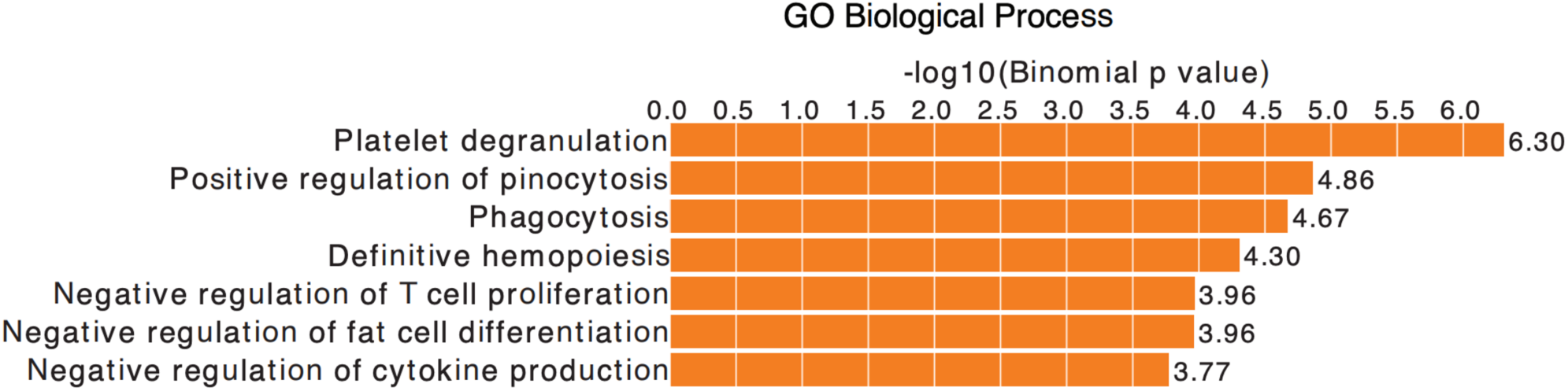
Gene Ontology enrichment of cfPeaks absent in WBC chromatin. GREAT analysis of cell-free cQTLs peaks (N = 766) present specifically in cell-free H3K27ac chromatin peaks but not in immune-cell H3K27ac chromatin peaks identified significant enrichment in both Gene Ontology Biological Process terms.

**Supplementary Fig. 4.**
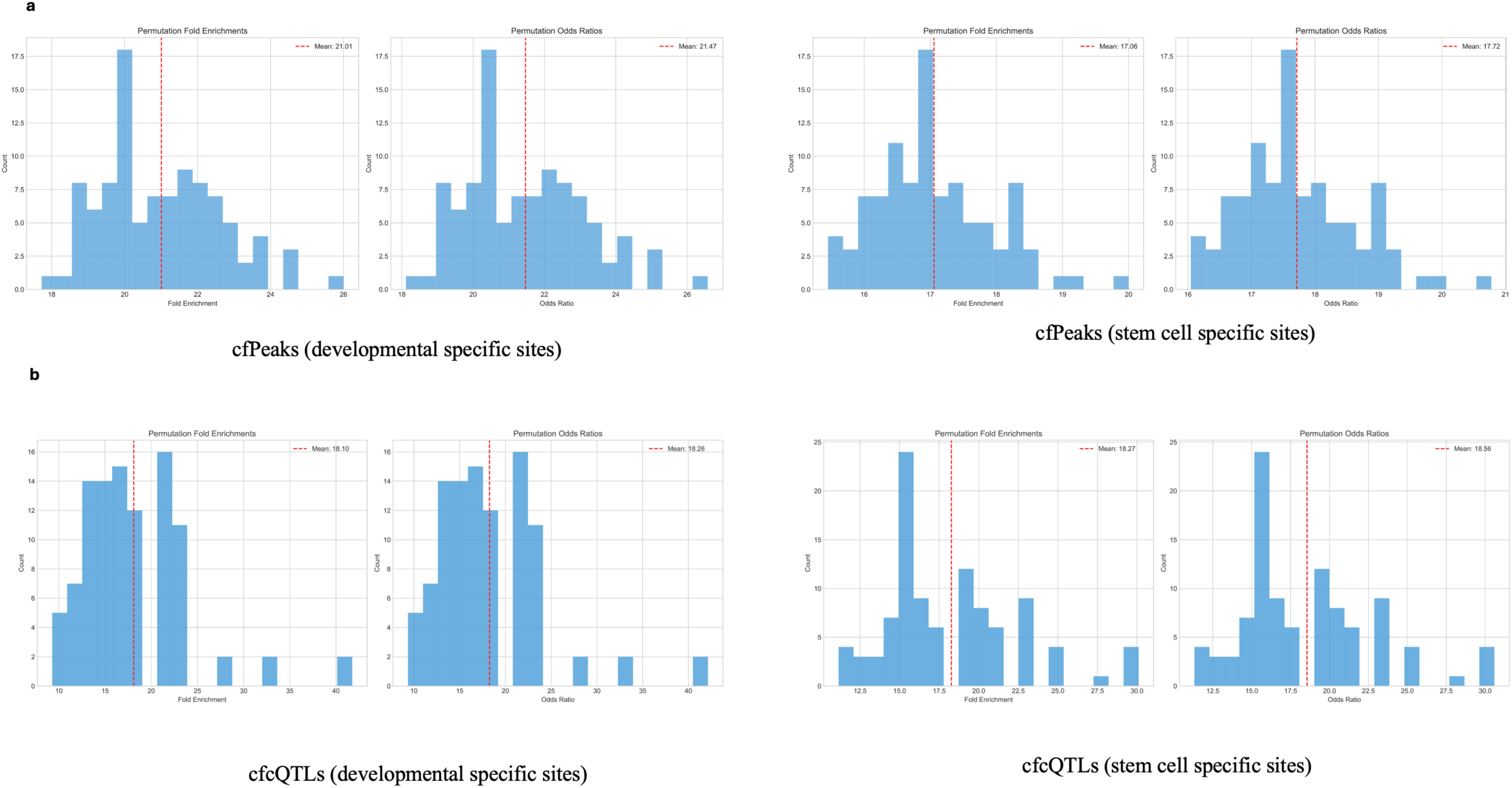
Permutation-mean fold enrichment. **(a)** cfPeaks enrichment at “active” associated specific sites compared to WBC peaks for developmental specific sites (left) and stem cell specific sites (right) **(b)** cfcQTLs enrichment at “active” associated specific sites compared to WBC-cQTLs for developmental specific sites (left) and stem cell specific sites (right)

**Supplementary Fig. 5.**
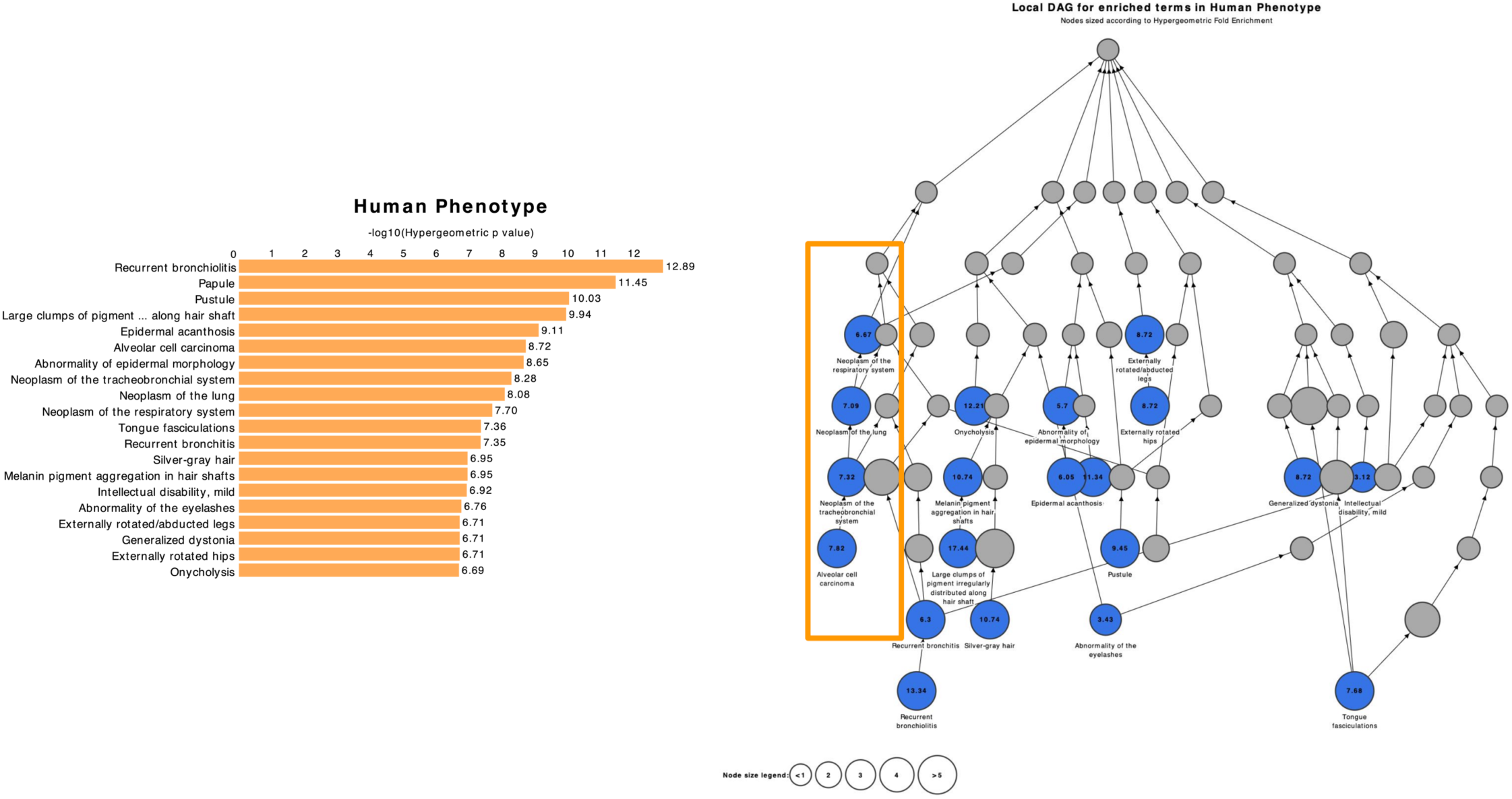
Human Phenotype enrichment network for cfcQTLs overlapping ctDNA-correlated regulatory elements. Human Phenotype enrichment analysis of cfcQTLs overlapping ctDNA-correlated regulatory elements (left panel). Local directed acyclic graph (DAG) showing enriched Human Phenotype terms with nodes sized according to hypergeometric fold enrichment. Blue nodes represent significantly enriched phenotypes with -log₁₀ P values indicated. Orange box highlights respiratory system neoplasm cluster including tracheobronchial system neoplasms, lung neoplasms, and respiratory system carcinomas (right panel)

**Supplementary Fig. 6.**
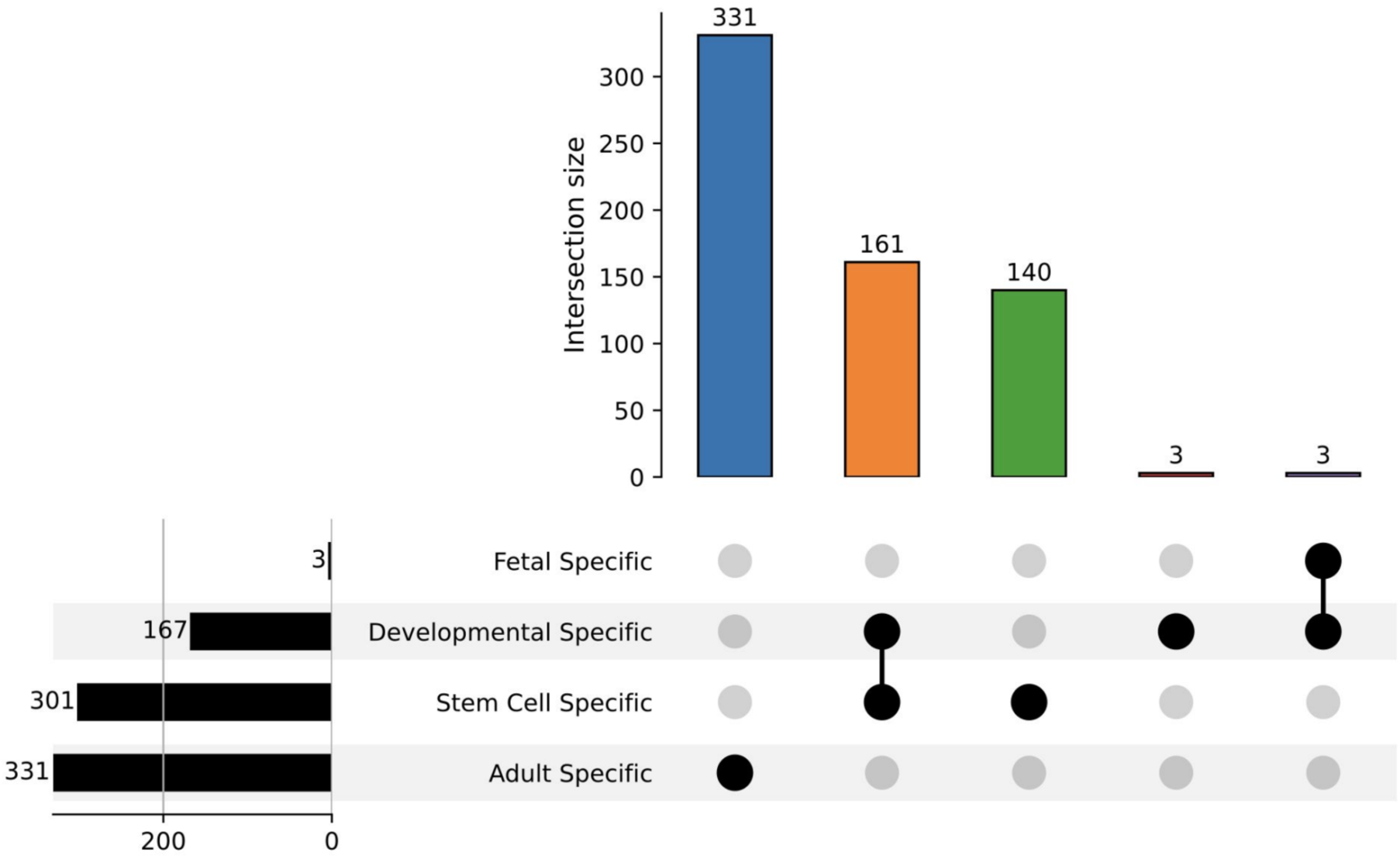
UpSet plot analysis of cfcQTL overlap across developmental specificity categories. UpSet plot showing intersection patterns of cfcQTLs across developmental categories. Most cfcQTLs are specific to single categories: 331 adult-specific only, 161 developmental-specific only, 140 stem cell-specific only, and 3 fetal-specific only. Minimal overlap between categories indicates distinct regulatory landscapes captured by each developmental stage

**Supplementary Fig. 7.**
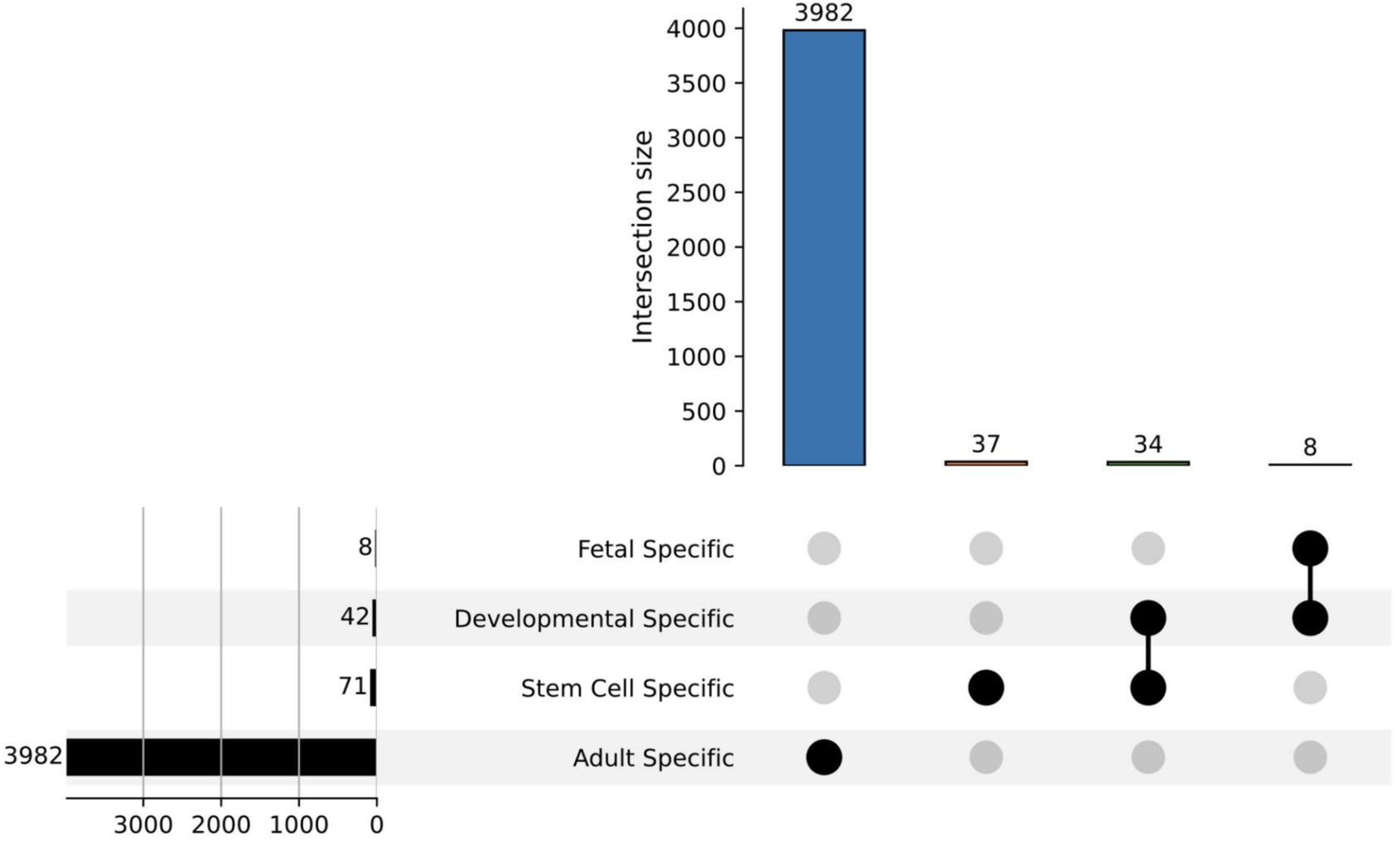
UpSet plot analysis of WBC-cQTL overlap across developmental specificity categories. UpSet plot showing intersection patterns of WBC-cQTLs across developmental categories. WBC-cQTLs show overwhelming bias toward adult-specific regulatory elements (3,982 out of total) with minimal representation in developmental contexts: 71 stem cell-specific, 42 developmental-specific, and 8 fetal-specific, showing restricted capture of non-adult regulatory programs.

**Supplementary Fig. 8.**
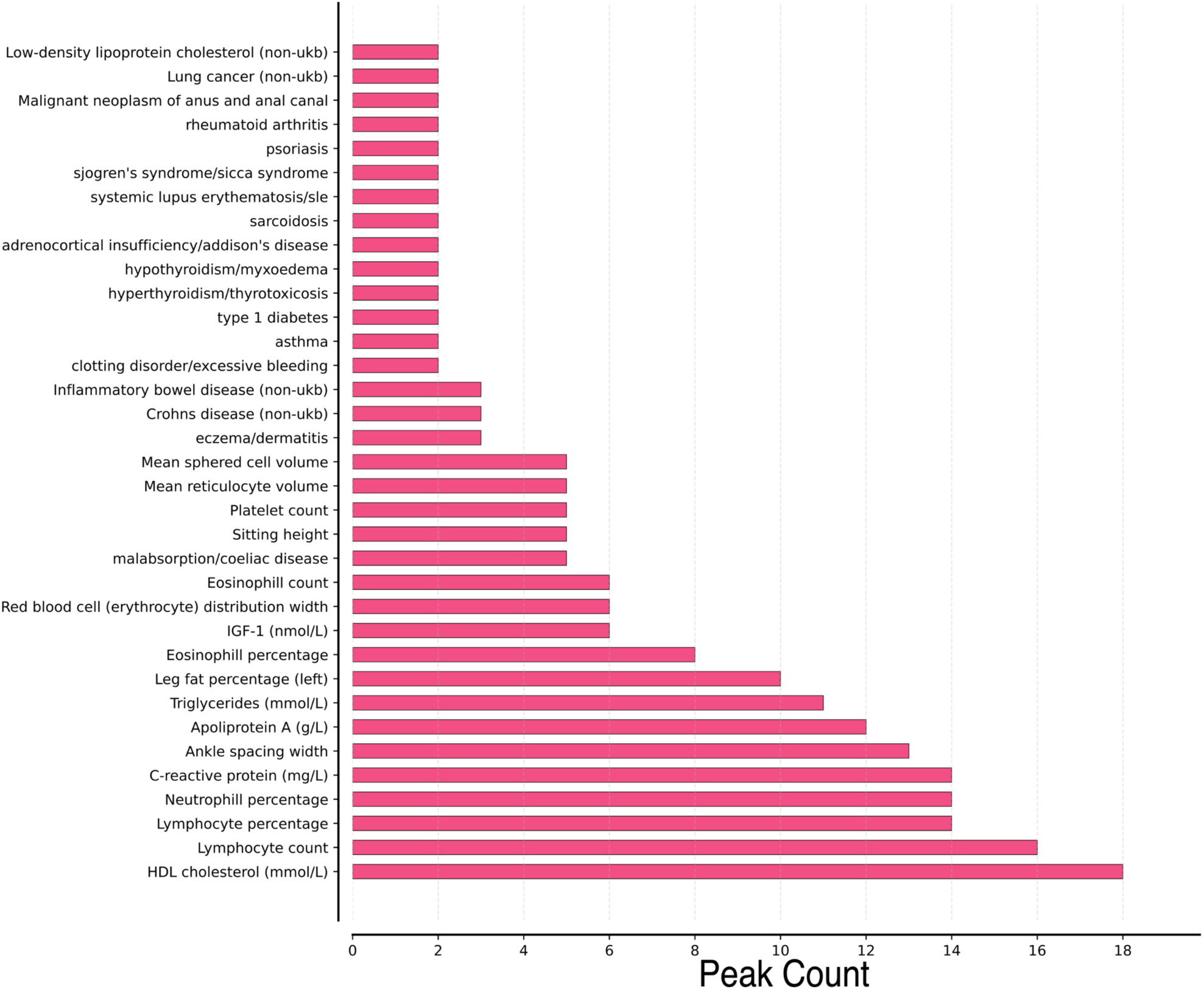
CWAS associations for adult-specific private cfcQTLs. Top 35 trait associations for private cfcQTLs overlapping adult-specific regulatory elements, ranked by peak count. Associations include metabolic biomarkers (HDL cholesterol, triglycerides), hematological traits (lymphocyte and eosinophil measurements), autoimmune diseases (rheumatoid arthritis, inflammatory bowel disease, Crohn’s disease, psoriasis, Type 1 diabetes), and cancer phenotypes (lung cancer). These associations were completely absent in private WBC-cQTLs.

**Supplementary Fig. 9.**
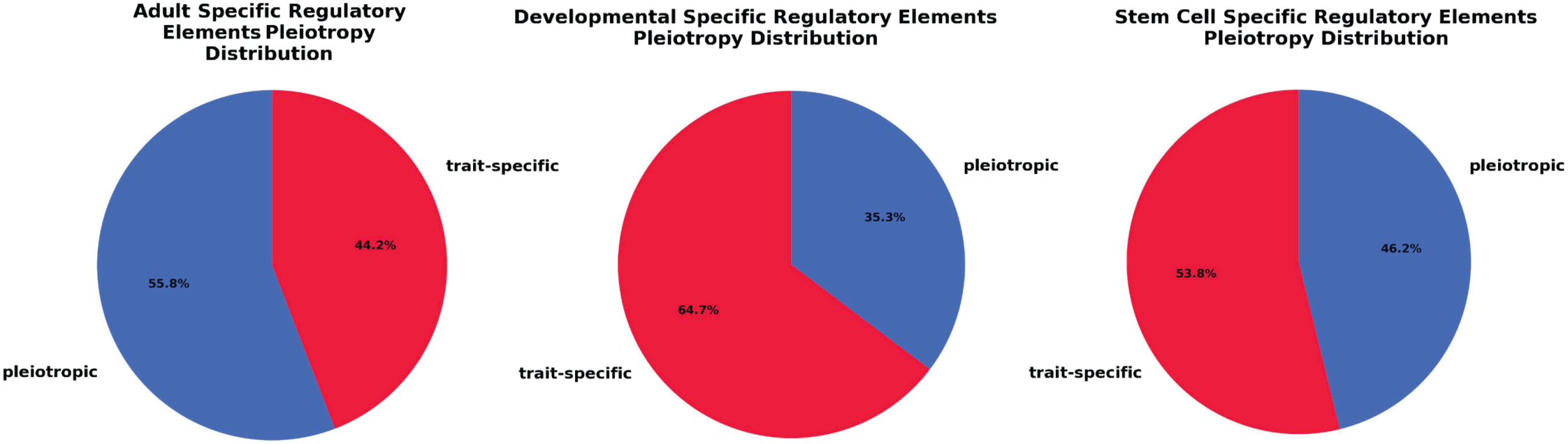
Distribution of trait associations across developmental categories. Pie charts showing trait-specific versus multi-trait associations for private cfcQTLs in adult-specific, developmental-specific, and stem cell-specific regulatory elements. Developmental-specific elements show more trait specificity (64.7%), compared to adult-specific (44.2%).

**Supplementary Fig. 10.**
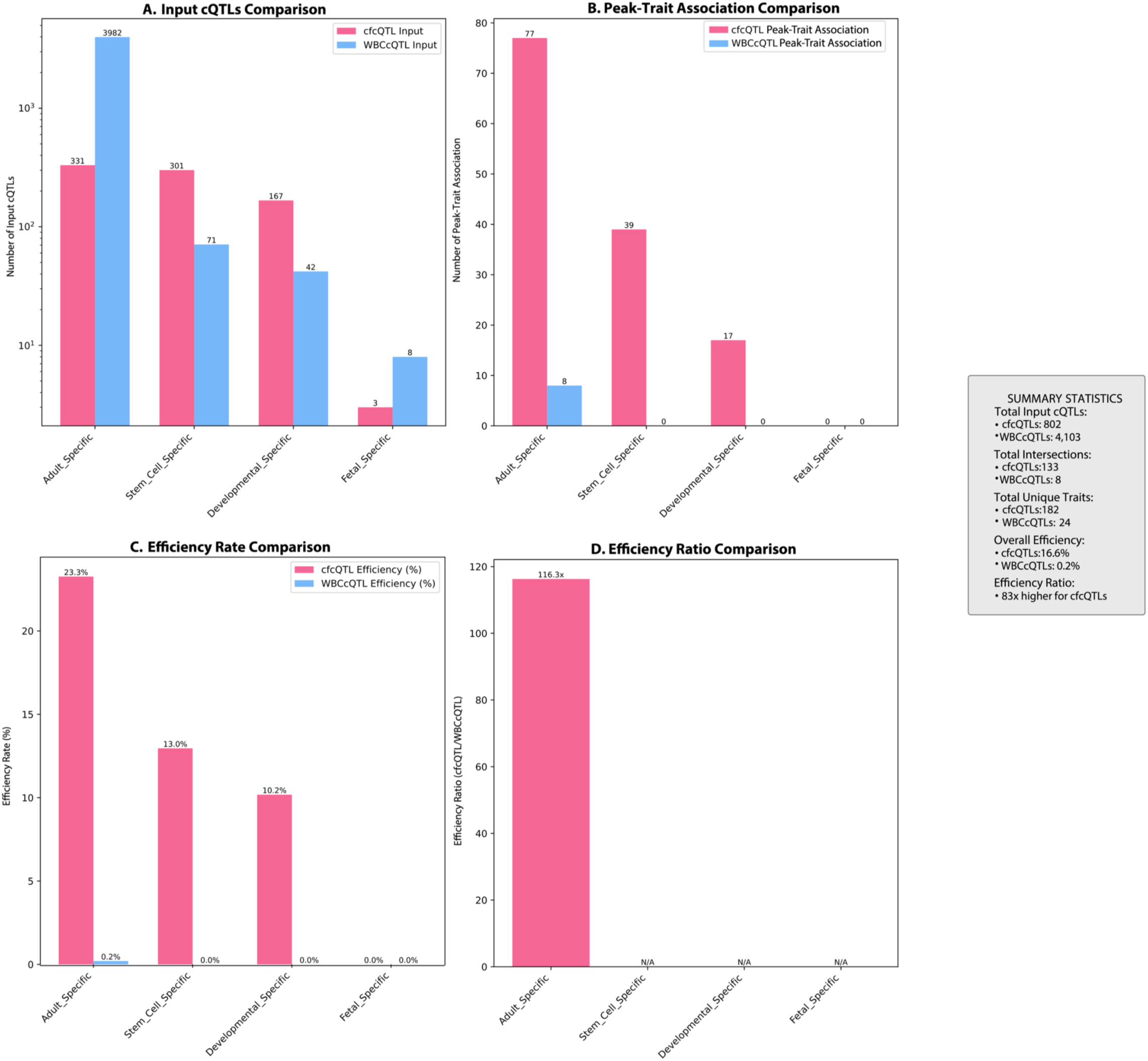
Regulatory efficiency comparison between private cfcQTLs and private WBC-cQTLs across developmental categories. **(a)** Input cQTL counts for private cfcQTLs (pink) and private WBC-cQTLs (blue) stratified by developmental stage (adult-specific, stem cell-specific, developmental-specific, fetal-specific). **(b)** Number of significant peak-trait associations identified through CWAS for each developmental category. **(c)** Association efficiency rates (percentage of cQTLs yielding significant trait associations) for cfcQTLs and WBC-cQTLs **(d)** Efficiency ratio showing fold-difference in association rates between cfcQTLs and WBC-cQTLs.

**Extended Data Fig. 1.**
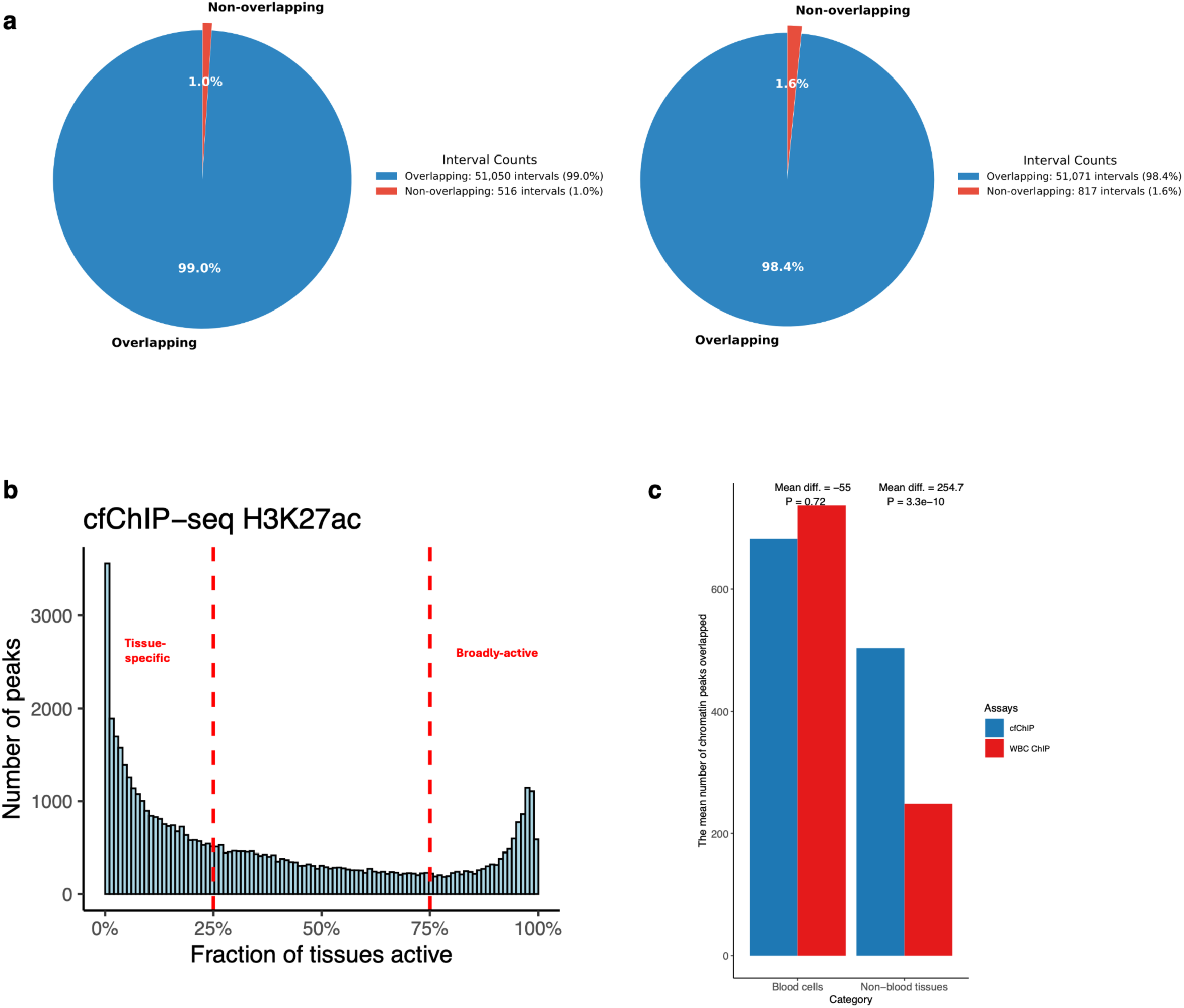
Tissue specificity analysis of cfChIP-seq regulatory elements. **(a)** Direct validation of cfPeaks with Roadmap Epigenomics tissue datasets. Pie charts show the proportion of cfPeaks that directly overlap with corresponding histone modification peaks from all Roadmap tissues. H3K27ac cfPeaks show 99.0% overlap (left) and H3K4me3 cfPeaks show 98.4% overlap (right) with tissue-derived ChIP-seq peaks, confirming that cfPeaks represent true regulatory elements. **(b)** Distribution of cfChIP-seq H3K27ac peaks by tissue activity breadth across 98 Roadmap reference epigenomes. Peaks are binned by the fraction of tissues in which they are active. Red dashed lines delineate tissue-specific peaks (active in <25% of tissues) from broadly-active peaks (active in >75% of tissues). The majority of cfPeaks show restricted tissue activity patterns. **(c)** Comparison of tissue-specific regulatory element capture between cfChIP-seq and WBC ChIP-seq for peaks active in <25% of tissues across blood cell types versus non-blood tissues. Bars represent mean number of chromatin peaks overlapping with reference epigenomes, stratified by tissue category. Statistical comparisons performed using Wilcoxon rank test. cfChIP-seq shows similar capture of blood cell regulatory elements but significantly greater overlap with non-blood tissue elements compared to WBC ChIP-seq.

**Extended Data Fig. 2.**
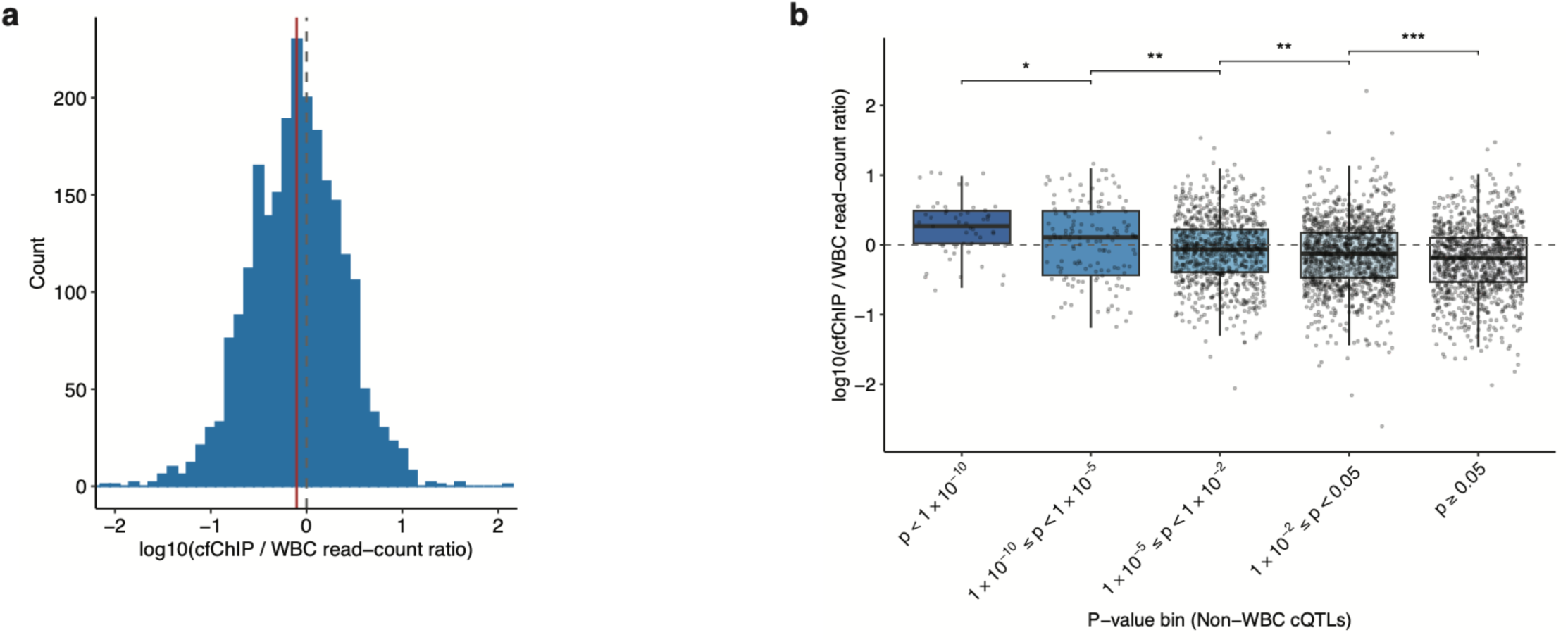
Cell-free cQTLs reveal context-dependent effects on gene regulation. **(a)** Distribution of log_₁₀_(cfChIP/WBC read count ratios) for cfChIP-specific allelic imbalance regions. **(b)** cfChIP/WBC read count ratios (log₁₀ scale, y-axis) stratified by allelic imbalance significance levels in cfChIP-seq. Higher significance regions show elevated cfChIP coverage relative to WBC ChIP-seq (*P < 0.05, ** P < 0.01, ***P < 0.001).

**Extended Data Fig. 3.**
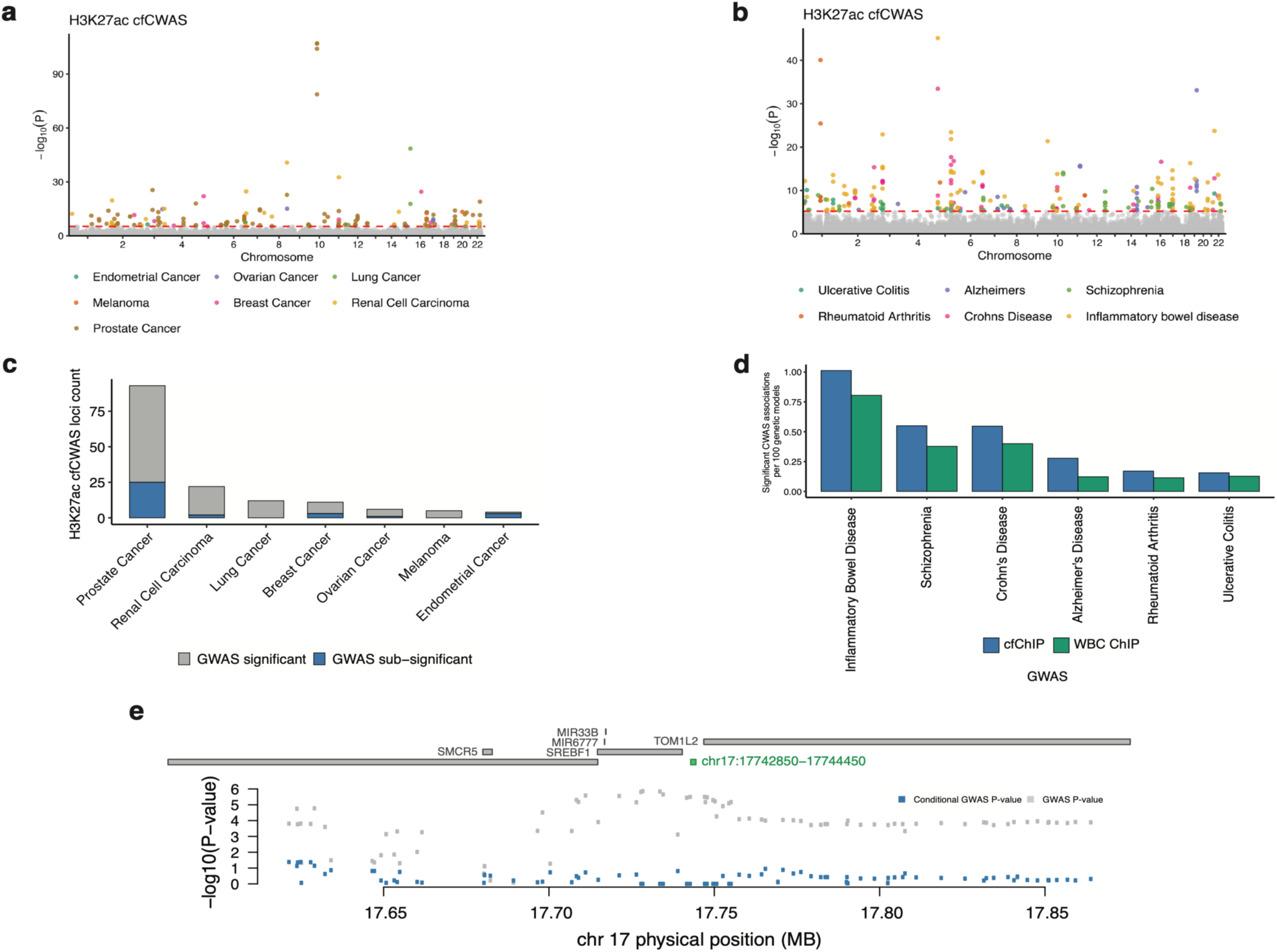
CWAS identifies novel cancer and non-cancer risk loci using cell-free epigenomic reference panels. **(a,b)** Manhattan plots depicting genome-wide significant associations (−log₁₀(P)) for multiple **(a)** cancer and **(b)**non-cancer types using H3K27ac cfChIP-seq reference panels, with colors denoting different cancer types as indicated in the figure legend. **(c)** Analysis of loci with significant H3K27ac cfCWAS associations, distinguishing between GWAS-significant loci (gray) and sub-threshold GWAS loci (blue) that achieved statistical significance only through the enhanced statistical power of the CWAS framework **(d)** Quantitative comparison of CWAS performance using cfChIP H3K27ac versus WBC ChIP H3K27ac reference panels across six non-cancer types. The y-axis represents the percentage of genetically determined chromatin accessibility regions harboring significant disease associations, illustrating consistently elevated detection rates with cfChIP-derived reference panels (blue) compared to WBC-derived references (green). **(e)** Regional association plot of chromosome 17q11.2 demonstrating attenuation of prostate cancer GWAS signal after conditioning on H3K27ac cfCWAS signal.

**Extended Data Fig. 5.**
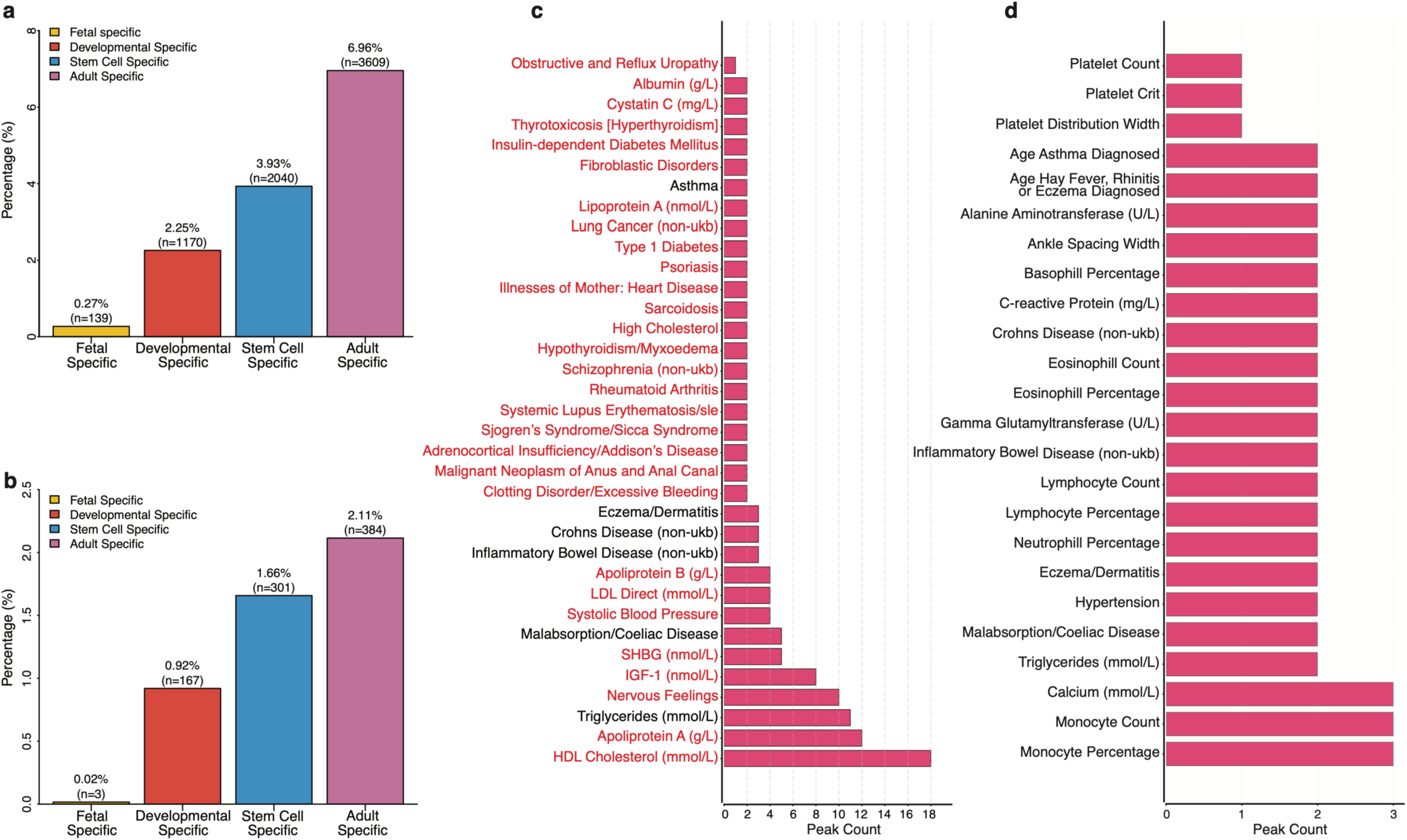
Detailed analysis of cfcQTL enrichment at developmental regulatory elements and disease associations. **(a)** Percentage distribution of cfPeaks across developmental-stage regulatory elements. Adult-specific sites comprise 6.96% (n=3,614), stem cell-specific sites comprise 3.93% (n=2,040), developmental-specific sites comprise 2.25% (n=1,170), and fetal-specific sites comprise 0.27% (n=139). **(b)** Percentage distribution of H3K27ac cfcQTLs across developmental-stage regulatory elements. Adult-specific sites comprise 2.11% (n=384), stem cell-specific sites comprise 1.66% (n=301), developmental-specific sites comprise 0.92% (n=167), and fetal-specific sites comprise minimal representation (n=3). (c) Traits associated with private cfcQTLs across developmentally stratified regulatory regions (n=35 traits), highlighting metabolic, immune, and disease-relevant phenotypes. Red indicates traits captured exclusively by private cfcQTLs but not by private WBC-cQTLs. (d) Traits associated with private WBC-cQTLs across developmentally stratified regulatory regions (n=24 traits).

## References

1. Maurano, M. T. et al. Systematic localization of common disease-associated variation in regulatory DNA. Science 337, 1190–1195 (2012).

2. Albert, F. W. & Kruglyak, L. The role of regulatory variation in complex traits and disease. Nat. Rev. Genet. 16, 197–212 (2015).

3. Gusev, A. et al. Partitioning heritability of regulatory and cell-type-specific variants across 11 common diseases. Am. J. Hum. Genet. 95, 535–552 (2014).

4. Visscher, P. M. et al. 10 years of GWAS discovery: Biology, function, and translation. Am. J. Hum. Genet. 101, 5–22 (2017).

5. Gallagher, M. D. & Chen-Plotkin, A. S. The post-GWAS era: From association to function. Am. J. Hum. Genet. 102, 717–730 (2018).

6. Buniello, A. et al. The NHGRI-EBI GWAS Catalog of published genome-wide association studies, targeted arrays and summary statistics 2019. Nucleic Acids Res. 47, D1005–D1012 (2019).

7. Alasoo, K. et al. Shared genetic effects on chromatin and gene expression indicate a role for enhancer priming in immune response. Nat. Genet. 50, 424–431 (2018).

8. Baca, S. C. et al. Genetic determinants of chromatin reveal prostate cancer risk mediated by context-dependent gene regulation. Nat. Genet. 54, 1364–1375 (2022).

9. Strober, B. J. et al. Dynamic genetic regulation of gene expression during cellular differentiation. Science 364, 1287–1290 (2019).

10. Mu, Z. et al. Impact of disease-associated chromatin accessibility QTLs across immune cell types and contexts. medRxiv (2024) doi:10.1101/2024.12.05.24318552.

11. GTEx Consortium. The GTEx Consortium atlas of genetic regulatory effects across human tissues. Science 369, 1318–1330 (2020).

12. Aguet, F. et al. Molecular quantitative trait loci. Nat. Rev. Methods Primers 3, (2023).

13. Umans, B. D., Battle, A. & Gilad, Y. Where are the disease-associated eQTLs? Trends Genet. 37, 109–124 (2021).

14. Kim-Hellmuth, S. et al. Genetic regulatory effects modified by immune activation contribute to autoimmune disease associations. Nat. Commun. 8, 266 (2017).

15. Soskic, B. et al. Immune disease risk variants regulate gene expression dynamics during CD4+ T cell activation. Nat. Genet. 54, 817–826 (2022).

16. Fairfax, B. P. et al. Innate immune activity conditions the effect of regulatory variants upon monocyte gene expression. Science 343, 1246949 (2014).

17. Yao, D. W., O’Connor, L. J., Price, A. L. & Gusev, A. Quantifying genetic effects on disease mediated by assayed gene expression levels. Nat. Genet. 52, 626–633 (2020).

18. Mu, Z. et al. The impact of cell type and context-dependent regulatory variants on human immune traits. Genome Biol. 22, 122 (2021).

19. Degner, J. F. et al. DNase I sensitivity QTLs are a major determinant of human expression variation. Nature 482, 390–394 (2012).

20. McVicker, G. et al. Identification of genetic variants that affect histone modifications in human cells. Science 342, 747–749 (2013).

21. Jeong, R. & Bulyk, M. L. Chromatin accessibility variation provides insights into missing regulation underlying immune-mediated diseases. bioRxivorg 2024.04.12.589213 (2024) doi:10.1101/2024.04.12.589213.

22. Arthur, T. D. et al. Multiomic QTL mapping reveals phenotypic complexity of GWAS loci and prioritizes putative causal variants. Cell Genom. 5, 100775 (2025).

23. Aracena, K. A. et al. Epigenetic variation impacts individual differences in the transcriptional response to influenza infection. Nat. Genet. 56, 408–419 (2024).

24. Grishin, D. & Gusev, A. Allelic imbalance of chromatin accessibility in cancer identifies candidate causal risk variants and their mechanisms. Nat. Genet. 54, 837–849 (2022).

25. Yanchus, C. et al. A noncoding single-nucleotide polymorphism at 8q24 drives IDH1-mutant glioma formation. Science 378, 68–78 (2022).

26. Spisak, S. et al. A biallelic multiple nucleotide length polymorphism explains functional causality at 5p15.33 prostate cancer risk locus. Nat. Commun. 14, 5118 (2023).

27. Patel, S. A. et al. The renal lineage factor PAX8 controls oncogenic signalling in kidney cancer. Nature 606, 999–1006 (2022).

28. Fulco, C. P. et al. Activity-by-contact model of enhancer-promoter regulation from thousands of CRISPR perturbations. Nat. Genet. 51, 1664–1669 (2019).

29. Mumbach, M. R. et al. HiChIP: efficient and sensitive analysis of protein-directed genome architecture. Nat. Methods 13, 919–922 (2016).

30. Armaos, A. et al. The PENGUIN approach to reconstruct protein interactions at enhancer-promoter regions and its application to prostate cancer. Nat. Commun. 14, 8084 (2023).

31. Benaglio, P. et al. Mapping genetic effects on cell type-specific chromatin accessibility and annotating complex immune trait variants using single nucleus ATAC-seq in peripheral blood. PLoS Genet. 19, e1010759 (2023).

32. Chen, L. et al. Genetic drivers of epigenetic and transcriptional variation in human immune cells. Cell 167, 1398–1414.e24 (2016).

33. Cristiano, S. et al. Genome-wide cell-free DNA fragmentation in patients with cancer. Nature 570, 385–389 (2019).

34. Snyder, M. W., Kircher, M., Hill, A. J., Daza, R. M. & Shendure, J. Cell-free DNA comprises an in vivo nucleosome footprint that informs its tissues-of-origin. Cell 164, 57–68 (2016).

35. Lo, Y. M. D. et al. Maternal plasma DNA sequencing reveals the genome-wide genetic and mutational profile of the fetus. Sci. Transl. Med. 2, 61ra91 (2010).

36. Wan, J. C. M. et al. Liquid biopsies come of age: towards implementation of circulating tumour DNA. Nat. Rev. Cancer 17, 223–238 (2017).

37. Adalsteinsson, V. A. et al. Scalable whole-exome sequencing of cell-free DNA reveals high concordance with metastatic tumors. Nat. Commun. 8, (2017).

38. Truszewska, A., Foroncewicz, B. & Pączek, L. The role and diagnostic value of cell-free DNA in systemic lupus erythematosus. Clin. Exp. Rheumatol. 35, 330–336 (2017).

39. Duvvuri, B. & Lood, C. Cell-free DNA as a biomarker in autoimmune rheumatic diseases. Front. Immunol. 10, 502 (2019).

40. Rykova, E. et al. Circulating DNA in rheumatoid arthritis: pathological changes and association with clinically used serological markers. Arthritis Res. Ther. 19, 85 (2017).

41. Rainer, T. H. et al. Prognostic use of circulating plasma nucleic acid concentrations in patients with acute stroke. Clin. Chem. 49, 562–569 (2003).

42. Zemmour, H. et al. Non-invasive detection of human cardiomyocyte death using methylation patterns of circulating DNA. Nat. Commun. 9, 1443 (2018).

43. Khemka, S., Sehar, U., Manna, P. R., Kshirsagar, S. & Reddy, P. H. Cell-free DNA as peripheral biomarker of Alzheimer’s disease. Aging Dis. 16, 787–803 (2024).

44. Gaitsch, H., Franklin, R. J. M. & Reich, D. S. Cell-free DNA-based liquid biopsies in neurology. Brain 146, 1758–1774 (2023).

45. Dwivedi, D. J. et al. Prognostic utility and characterization of cell-free DNA in patients with severe sepsis. Crit. Care 16, R151 (2012).

46. Norton, M. E. et al. Cell-free DNA analysis for noninvasive examination of trisomy. N. Engl. J. Med. 372, 1589–1597 (2015).

47. Sadeh, R. et al. ChIP-seq of plasma cell-free nucleosomes identifies gene expression programs of the cells of origin. Nat. Biotechnol. 39, 586–598 (2021).

48. Baca, S. C. et al. Liquid biopsy epigenomic profiling for cancer subtyping. Nat. Med. 29, 2737–2741 (2023).

49. Roadmap Epigenomics Consortium et al. Integrative analysis of 111 reference human epigenomes. Nature 518, 317–330 (2015).

50. Stunnenberg, H. G. et al. The international human epigenome consortium: A blueprint for scientific collaboration and discovery. Cell 167, 1145–1149 (2016).

51. Choi, B. J. et al. Altered expression of CDX2 in colorectal cancers. APMIS 114, 50–54 (2006).

52. Baba, Y. et al. Relationship of CDX2 loss with molecular features and prognosis in colorectal cancer. Clin. Cancer Res. 15, 4665–4673 (2009).

53. Badia-Ramentol, J. et al. The prognostic potential of CDX2 in colorectal cancer: Harmonizing biology and clinical practice. Cancer Treat. Rev. 121, 102643 (2023).

54. Guo, Z., Wang, Y., Xiang, S., Wang, S. & Chan, F. L. Chromogranin A is a predictor of prognosis in patients with prostate cancer: a systematic review and meta-analysis. Cancer Manag. Res. 11, 2747–2758 (2019).

55. Ewing, C. M. et al. Germline mutations in HOXB13 and prostate-cancer risk. N. Engl. J. Med. 366, 141–149 (2012).

56. Darst, B. F. et al. A rare germline HOXB13 variant contributes to risk of prostate cancer in men of African ancestry. Eur. Urol. 81, 458–462 (2022).

57. Brechka, H., Bhanvadia, R. R., VanOpstall, C. & Vander Griend, D. J. HOXB13 mutations and binding partners in prostate development and cancer: Function, clinical significance, and future directions. Genes Dis. 4, 75–87 (2017).

58. Al Olama, A. A. et al. A meta-analysis of 87,040 individuals identifies 23 new susceptibility loci for prostate cancer. Nat. Genet. 46, 1103–1109 (2014).

59. Amin Al Olama, A., et al. Multiple novel prostate cancer susceptibility signals identified by fine-mapping of known risk loci among Europeans. Hum. Mol. Genet. 24, 5589–5602 (2015).

60. Schumacher, F. R. et al. Genome-wide association study identifies new prostate cancer susceptibility loci. Hum. Mol. Genet. 20, 3867–3875 (2011).

61. Eeles, R. A. et al. Multiple newly identified loci associated with prostate cancer susceptibility. Nat. Genet. 40, 316–321 (2008).

62. Kote-Jarai, Z. et al. Fine-mapping identifies multiple prostate cancer risk loci at 5p15, one of which associates with TERT expression. Hum. Mol. Genet. 22, 2520–2528 (2013).

63. Chen, H. et al. Large-scale cross-cancer fine-mapping of the 5p15.33 region reveals multiple independent signals. HGG Adv. 2, 100041 (2021).

64. Grampp, S. et al. Genetic variation at the 8q24.21 renal cancer susceptibility locus affects HIF binding to a MYC enhancer. Nat. Commun. 7, 13183 (2016).

65. Shetty, A. et al. Allele-specific epigenetic activity in prostate cancer and normal prostate tissue implicates prostate cancer risk mechanisms. Am. J. Hum. Genet. 108, 2071–2085 (2021).

66. Delaneau, O. et al. A complete tool set for molecular QTL discovery and analysis. Nat. Commun. 8, 15452 (2017).

67. Kellman, L. N. et al. Functional analysis of cancer-associated germline risk variants. Nat. Genet. 57, 718–728 (2025).

68. Cheng, T. H. T. et al. Meta-analysis of genome-wide association studies identifies common susceptibility polymorphisms for colorectal and endometrial cancer near SH2B3 and TSHZ1. Sci. Rep. 5, 17369 (2015).

69. McKay, J. D. et al. Large-scale association analysis identifies new lung cancer susceptibility loci and heterogeneity in genetic susceptibility across histological subtypes. Nat. Genet. 49, 1126–1132 (2017).

70. Wu, C. et al. Genome-wide association analyses of esophageal squamous cell carcinoma in Chinese identify multiple susceptibility loci and gene-environment interactions. Nat. Genet. 44, 1090–1097 (2012).

71. Michailidou, K. et al. Association analysis identifies 65 new breast cancer risk loci. Nature 551, 92–94 (2017).

72. Yoon, K.-A. et al. A genome-wide association study reveals susceptibility variants for non-small cell lung cancer in the Korean population. Hum. Mol. Genet. 19, 4948–4954 (2010).

73. Melin, B. S. et al. Genome-wide association study of glioma subtypes identifies specific differences in genetic susceptibility to glioblastoma and non-glioblastoma tumors. Nat. Genet. 49, 789–794 (2017).

74. Boix, C. A., James, B. T., Park, Y. P., Meuleman, W. & Kellis, M. Regulatory genomic circuitry of human disease loci by integrative epigenomics. Nature 590, 300–307 (2021).

75. Gusev, A., et al. Allelic imbalance reveals widespread germline-somatic regulatory differences and prioritizes risk loci in Renal Cell Carcinoma. bioRxiv 631150 (2019) doi:10.1101/631150.

76. Vorperian, S. K., Moufarrej, M. N., Tabula Sapiens Consortium & Quake, S. R. Cell types of origin of the cell-free transcriptome. Nat. Biotechnol. 40, 855–861 (2022).

77. Moss, J. et al. Megakaryocyte- and erythroblast-specific cell-free DNA patterns in plasma and platelets reflect thrombopoiesis and erythropoiesis levels. Nat. Commun. 14, 7542 (2023).

78. Kammers, K. et al. Transcriptional profile of platelets and iPSC-derived megakaryocytes from whole-genome and RNA sequencing. Blood 137, 959–968 (2021).

79. Zhang, H. et al. Genome-wide association study identifies 32 novel breast cancer susceptibility loci from overall and subtype-specific analyses. Nat. Genet. 52, 572–581 (2020).

80. Schumacher, F. R. et al. Association analyses of more than 140,000 men identify 63 new prostate cancer susceptibility loci. Nat. Genet. 50, 928–936 (2018).

81. Purdue, M. P. et al. Multi-ancestry genome-wide association study of kidney cancer identifies 63 susceptibility regions. Nat. Genet. 56, 809–818 (2024).

82. Gharahkhani, P. et al. Genome-wide association studies in oesophageal adenocarcinoma and Barrett’s oesophagus: a large-scale meta-analysis. Lancet Oncol. 17, 1363–1373 (2016).

83. Phelan, C. M. et al. Identification of 12 new susceptibility loci for different histotypes of epithelial ovarian cancer. Nat. Genet. 49, 680–691 (2017).

84. Rashkin, S. R. et al. Pan-cancer study detects genetic risk variants and shared genetic basis in two large cohorts. Nat. Commun. 11, 4423 (2020).

85. Huyghe, J. R. et al. Discovery of common and rare genetic risk variants for colorectal cancer. Nat. Genet. 51, 76–87 (2019).

86. Lee, J. C. et al. Genome-wide association study identifies distinct genetic contributions to prognosis and susceptibility in Crohn’s disease. Nat. Genet. 49, 262–268 (2017).

87. de Lange, K. M. et al. Genome-wide association study implicates immune activation of multiple integrin genes in inflammatory bowel disease. Nat. Genet. 49, 256–261 (2017).

88. Stahl, E. A. et al. Genome-wide association study meta-analysis identifies seven new rheumatoid arthritis risk loci. Nat. Genet. 42, 508–514 (2010).

89. McGovern, D. P. B. et al. Genome-wide association identifies multiple ulcerative colitis susceptibility loci. Nat. Genet. 42, 332–337 (2010).

90. Saklatvala, J. R. et al. Genetic validation of psoriasis phenotyping in UK biobank supports the utility of self-reported data and composite definitions for large genetic and epidemiological studies. J. Invest. Dermatol. 143, 1598–1601.e10 (2023).

91. Ripke, S. et al. Genome-wide association analysis identifies 13 new risk loci for schizophrenia. Nat. Genet. 45, 1150–1159 (2013).

92. Jansen, I. E. et al. Author Correction: Genome-wide meta-analysis identifies new loci and functional pathways influencing Alzheimer’s disease risk. Nat. Genet. 52, 354 (2020).

93. Wang, A. et al. Characterizing prostate cancer risk through multi-ancestry genome-wide discovery of 187 novel risk variants. Nat. Genet. 55, 2065–2074 (2023).

94. Sáez, C. et al. Expression of basic fibroblast growth factor and its receptors FGFR1 and FGFR2 in human benign prostatic hyperplasia treated with finasteride. Prostate 40, 83–88 (1999).

95. Boget, S., Cereser, C., Parvaz, P., Leriche, A. & Revol, A. Fibroblast growth factor receptor 1 (FGFR1) is over-expressed in benign prostatic hyperplasia whereas FGFR2-IIIc and FGFR3 are not. Eur. J. Endocrinol. 145, 303–310 (2001).

96. Corn, P. G., Wang, F., McKeehan, W. L. & Navone, N. Targeting fibroblast growth factor pathways in prostate cancer. Clin. Cancer Res. 19, 5856–5866 (2013).

97. Wei, G. et al. SREBF1-based metabolic reprogramming in prostate cancer promotes tumor ferroptosis resistance. Cell Death Discov. 11, 75 (2025).

98. Audet-Walsh, É. et al. SREBF1 activity is regulated by an AR/mTOR nuclear axis in prostate cancer. Mol. Cancer Res. 16, 1396–1405 (2018).

99. Bycroft, C. et al. The UK Biobank resource with deep phenotyping and genomic data. Nature 562, 203–209 (2018).

100. Pomerantz, M. M. et al. Prostate cancer reactivates developmental epigenomic programs during metastatic progression. Nat. Genet. 52, 790–799 (2020).

101. Thiery, J. P., Acloque, H., Huang, R. Y. J. & Nieto, M. A. Epithelial-mesenchymal transitions in development and disease. Cell 139, 871–890 (2009).

102. Tam, W. L. & Weinberg, R. A. The epigenetics of epithelial-mesenchymal plasticity in cancer. Nat. Med. 19, 1438–1449 (2013).

103. Polyak, K. & Weinberg, R. A. Transitions between epithelial and mesenchymal states: acquisition of malignant and stem cell traits. Nat. Rev. Cancer 9, 265–273 (2009).

104. Dietlein, F. et al. Genome-wide analysis of somatic noncoding mutation patterns in cancer. Science 376, eabg5601 (2022).

105. Rheinbay, E. et al. Analyses of non-coding somatic drivers in 2,658 cancer whole genomes. Nature 578, 102–111 (2020).

106. ICGC/TCGA Pan-Cancer Analysis of Whole Genomes Consortium. Pan-cancer analysis of whole genomes. Nature 578, 82–93 (2020).

107. Martincorena, I. & Campbell, P. J. Somatic mutation in cancer and normal cells. Science 349, 1483–1489 (2015).

108. Huang, F. W. et al. Highly recurrent TERT promoter mutations in human melanoma. Science 339, 957–959 (2013).

109. Killela, P. J. et al. TERT promoter mutations occur frequently in gliomas and a subset of tumors derived from cells with low rates of self-renewal. Proc. Natl. Acad. Sci. U. S. A. 110, 6021–6026 (2013).

110. Horn, S. et al. TERT promoter mutations in familial and sporadic melanoma. Science 339, 959–961 (2013).

111. Mostafavi, H., Spence, J. P., Naqvi, S. & Pritchard, J. K. Systematic differences in discovery of genetic effects on gene expression and complex traits. Nat. Genet. 55, 1866–1875 (2023).

112. Sun, T. et al. Systematic evaluation of methylation-based cell type deconvolution methods for plasma cell-free DNA. Genome Biol. 25, 318 (2024).

113. Sun, K. et al. Orientation-aware plasma cell-free DNA fragmentation analysis in open chromatin regions informs tissue of origin. Genome Res. 29, 418–427 (2019).

114. McCarthy, S. et al. A reference panel of 64,976 haplotypes for genotype imputation. Nat. Genet. 48, 1279–1283 (2016).

115. Davies, R. W., Flint, J., Myers, S. & Mott, R. Rapid genotype imputation from sequence without reference panels. Nat. Genet. 48, 965–969 (2016).

116. 1000 Genomes Project Consortium et al. A global reference for human genetic variation. Nature 526, 68–74 (2015).

117. Loh, P.-R. et al. Reference-based phasing using the Haplotype Reference Consortium panel. Nat. Genet. 48, 1443–1448 (2016).

118. van de Geijn, B., McVicker, G., Gilad, Y. & Pritchard, J. K. WASP: allele-specific software for robust molecular quantitative trait locus discovery. Nat. Methods 12, 1061–1063 (2015).

119. Castel, S. E., Levy-Moonshine, A., Mohammadi, P., Banks, E. & Lappalainen, T. Tools and best practices for data processing in allelic expression analysis. Genome Biol. 16, 195 (2015).

120. Kuilman, T. et al. CopywriteR: DNA copy number detection from off-target sequence data. Genome Biol. 16, 49 (2015).

121. Whitlock, M. C. Combining probability from independent tests: the weighted Z-method is superior to Fisher’s approach: Combining probabilities from many tests. J. Evol. Biol. 18, 1368–1373 (2005).

122. Gusev, A. et al. Integrative approaches for large-scale transcriptome-wide association studies. Nat. Genet. 48, 245–252 (2016).

123. McLean, C. Y. et al. GREAT improves functional interpretation of cis-regulatory regions. Nat. Biotechnol. 28, 495–501 (2010).

124. Quinlan, A. R. & Hall, I. M. BEDTools: a flexible suite of utilities for comparing genomic features. Bioinformatics 26, 841–842 (2010).

125. Dale, R. K., Pedersen, B. S. & Quinlan, A. R. Pybedtools: a flexible Python library for manipulating genomic datasets and annotations. Bioinformatics 27, 3423–3424 (2011).

